# KIASORT: Knowledge-Integrated Automated Spike Sorting for Geometry-Free Neuron Tracking

**DOI:** 10.1101/2025.07.10.664175

**Authors:** Kianoush Banaie Boroujeni, Thilo Womelsdorf, Sabine Kastner

**Affiliations:** Princeton Neuroscience Institute, Princeton University, Princeton, NJ 08544; Department of Psychology, Vanderbilt University, Nashville, TN 37212; Department of Psychology, Princeton University, Princeton, NJ 08544

## Abstract

Modern high-density neural recordings demand spike sorting algorithms that can handle diverse probe geometries and complex, neuron-specific drift, yet existing methods often rely on rigid geometric assumptions and one-dimensional drift models. Here, we introduce KIASORT (Knowledge-Integrated Automated Spike Sorting), a geometry-free approach for per-neuron drift tracking. KIASORT trains channel-specific classifiers in a hybrid linear–nonlinear embedding space, capturing waveform features often missed by conventional linear methods. These classifiers then sort spikes by independently tracking each neuron, unconstrained by probe layout. Biophysical simulations showed that even sub-micron probe displacements induce neuron-specific waveform distortions that standard drift models cannot correct. In ground-truth benchmarks with heterogeneous, neuron-specific drift, KIASORT significantly outperformed Kilosort4 in recovering high-quality units, while maintaining real-time performance on standard CPUs. Its robustness was further validated on both primate and mouse data. KIASORT combines automated sorting with manual curation in a unified graphical interface, offering a complete and user-friendly spike sorting platform. The software is freely available at https://kiasort.com.

## Introduction

Brain neural communication relies on the spiking activity of single neurons. While the spiking events are widely regarded as the smallest units of neural signaling, the accurate estimation and identification of the neuronal sources generating each spike from extracellular recordings remain a challenge. Such source identification is an essential step toward understanding neural circuit dynamics and compartmental computations in the brain ^1–3^. Over the past decades, the demand for distinguishing neuronal spiking activity in brain signals and reliably attributing it to its source has grown significantly with advances in electrophysiological recording technologies. In particular, high-density silicon probes such as Neuropixels, while enabling simultaneous recordings of large populations of neurons ^3,4^, impose an even greater need for robust and automated spike-sorting methods to reliably extract single-unit activity.

Traditionally, spike sorting involved three main steps: detection, feature extraction, and clustering ^5,6^. Early methods mostly relied on manual clustering of low dimensional features of spike waveforms, such as principal component analysis (PCA), and further manual adjustments ^6,7^. These techniques however faced limitations with regard to scalability, variability in spiking activity, and subjective biases in adjustments ^2,8,9^.

To resolve these challenges, more automated spike-sorting algorithms have been introduced to minimize manual curation while maximizing sorting accuracy and consistency ^3,8,10,11^. These algorithms often use a template-matching paradigm and are more optimal for high-density laminar probes, showing significant improvements in computational efficiency, scalability, and clustering performance compared with traditional approaches. Some examples include Kilosort ^11,12^, SpyKING Circus ^13^, MountainSort ^8^, and IronClust ^3^. Notably, Kilosort, and in particular Kilosort4, has been shown to outperform earlier algorithms by taking advantage of both the use of graph-based clustering and drift correction ^12^. Notably, Kilosort4 includes drift tracking and correction using piecewise rigid probe registration.

There are, however, key challenges yet to be resolved. Despite advances in graph-based clustering and drift correction, algorithmic cores still rest on rigid biophysical and statistical assumptions that real recordings frequently violate. First, spikes are modeled as linear, low-rank superpositions of fixed templates embedded in Gaussian noise; yet extracellular waveforms depend nonlinearly on neuronal morphology, initiation site, electrode orientation, and tissue conductivity, giving rise to polarity inversions, amplitude-only drifts, and clipped artifacts beyond the reach of any low-rank linear model ^14,15^. Second, the k-nearest-neighbour graph is constructed in a Euclidean PCA space whose principal axes remain warped by residual common-mode fluctuations and by subtle amplitude shifts as neurons drift relative to the shank. These anisotropic distortions, when judged by linear distance metrics, can still fracture sparse units into filaments or cause dense units to bleed together ^13,16^. Finally, most methods, including Kilosort4, correct only piecewise rigid motion along the z-axis under a uniform ≤ 40 µm pitch assumption; oblique insertions, wide Utah arrays, or angled Neuropixels 2 shanks introduce shear and rotation that remain unmodeled ^3,12^. These mismatches between algorithmic assumptions and biological reality become more pronounced when using different probe geometries, target deeper or axonal compartments in different species, or conduct long-term recordings in behaving animals.

Here, we introduce KIASORT (Knowledge-Integrated Automated Spike Sorting), a novel algorithm designed to overcome key limitations in current spike-sorting methods. KIASORT automatically detects and filters out channels with poor signal quality, dynamically removes variable recording noise, and clusters data using an iterative density-based approach applied to a hybrid dimensionality reduction framework that combines principal component analysis (PCA) with nonlinear uniform manifold approximation and projection (UMAP) embeddings. We show through biophysical modeling of multicompartmental neurons that drift patterns not aligned with the probe shank, even if subtle, lead to significant waveform changes depending on the neuron model and electrode proximity that conventional methods would leave uncorrected. To account for this, the algorithm incorporates a geometry-free, neuron-based tracking system that trains classifiers independently on each recording channel. This per-neuron tracking approach enables the identification of neurons across multiple channels and supports the cross-channel transfer of detected spikes to their reference channel for drift correction without any prior assumptions on drift type. These features allow the classifiers to adapt to nonlinear waveform changes and eliminate assumptions about cluster geometry, linear waveform superposition, or rigid or piecewise probe-level motion.

We evaluated KIASORT using simulated ground-truth multichannel datasets that closely mimic real-world recording conditions, including realistic noise profiles adapted from actual recordings, neuron-specific drift, nonlinear waveform transitions, and waveform heterogeneity. The simulation results show that KIASORT outperforms the current state-of-the-art method, Kilosort4, across several performance metrics and under different drift conditions. Specifically, KIASORT achieved 5-10% higher sorting precision and identified 5–15% more high-quality single units (defined as units with an F1 score exceeding 0.75), particularly under conditions with pronounced neuron-specific drift and nonlinear waveform changes. Although GPU acceleration is supported, KIASORT is optimized to achieve speeds comparable to real-time processing benchmarks using only standard CPUs, making it accessible to users without high-end computational hardware.

In addition to its automated sorting capabilities, KIASORT includes a graphical user interface (GUI) that integrates both sorting and post-hoc curation within a unified platform. This setup enables data exploration, user-defined parameter adjustments, and manual curation and verification of sorting results (**Figure S1**, see tutorial on https://kiasort.com for a guided tour of the KIASORT GUI). This all-in-one platform is a key feature that sets KIASORT apart from previous spike-sorting methods, which typically lack such integrated functionality.

## Results

The algorithm consists of three main modules (**Figure 1**). The first module (**i**) uniformly samples data across time to create a representative subset. The second module (**ii**) performs clustering on the extracted spikes from the sampled data. For each channel, spike waveforms are clustered using a feature vector constructed by UMAP and PCA embeddings through iterative density-based clustering. After merging similar clusters, labels are updated, and an SVM classifier is trained for each class using its PCA features. A reference channel is assigned to each cluster based on the channel where the spike amplitude is maximal. The third module (**iii**) carries out spike sorting on the full dataset by dividing it into small batches and using the trained classifiers to predict the label of each detected spike. This module also handles relabeling and cross-channel transfer for spikes whose reference channel differs from the originally assigned channel. During batch processing, the algorithm records outputs including spike times, features, waveforms (optional), channels, and labels.

**Figure 1.**
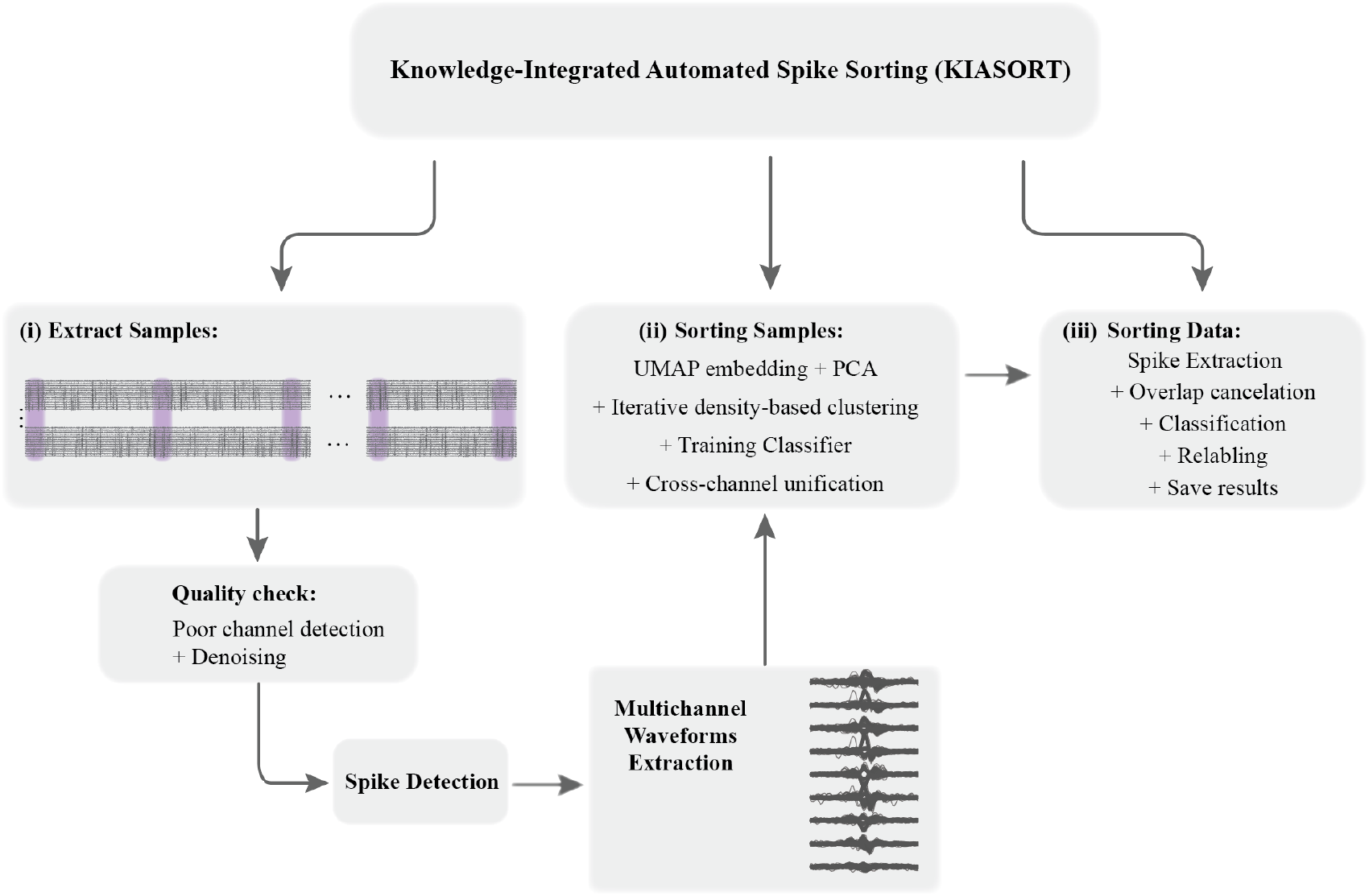
Main diagram of the KIASORT algorithm. The workflow consists of three modules for extracting and integrating knowledge for final classification of cells. **(i) Sample extraction:** Data are uniformly sampled across the recording to build a representative subset. During this step, channels are screened for poor-quality or silent electrodes, per-channel detection thresholds are defined, and multichannel spike waveforms are extracted for all threshold crossings. **(ii) Sample sorting:** Extracted waveforms are embedded using UMAP and PCA and concatenated into a joint feature space for iterative density-based clustering. Clusters that violate the refractory-period or ISI criteria are recursively split, highly correlated clusters are merged, labels are finalized, and an SVM classifier is trained on the PCA features of each unit. Each cluster is then assigned a reference channel based on the site of maximal absolute spike amplitude. **(iii) Data sorting:** The entire dataset is processed in small batches (with overlap cancellation). Spikes are detected and passed through the trained per-channel classifiers; spikes whose predicted reference-channel differs from the detection channel are transferred and retrieved at their reference-channel. Final outputs including spike times, features, waveforms (optional), channels, and unit labels are stored after processing each batch.

For the first module, initially, the algorithm performs a brief screening to detect poor-quality or silent channels. Each channel signal (in a short duration of 5 seconds) is decomposed into multiple frequency sub-bands and compares the power in the spike bands to low and high frequency sub-bands. We verified this method on simulated data with variable low and high frequency noise levels and spike band activities and show examples from real data recordings (**Figure S2**). Next, for denoising the signal, we used a two-pass percentile-template regression approach that builds a high-percentile noise template across channels and subtracts it adaptively from each channel. In low-SNR, high-channel-count recordings, signals can be dominated by slow common-mode drift on all or a subset of channels with moderate to high correlations (e.g., r ≈ 0.4-0.8). Even Kilosort4’s whitening, which uses spike-free covariance estimation, eigenvalue regularization, and is band-limited, can still over-whiten low-variance components which can distort spike shapes and smear high-frequency content in the signal. The percentile-template regression, however, only removes the empirically shared noise waveform at each time point, while preserving the residual spectrum and spike waveforms without ringing artifacts or attenuation of high-frequency spikes (see **Methods**).

Then a detection threshold is defined for each channel using median absolute deviation (MAD), and multichannel waveforms are extracted for all threshold-passing spikes. For each channel we store information about spikes such as spike indices and their multichannel waveforms. Note that each spike event will be detected at any given channel that passes the threshold. We only consider spikes that do not overlap with spikes on their own and nearby channels (see **Methods**).

### Per-channel clustering and classifier training

Once the spike samples are extracted, the second module (**Figure 1**) processes each channel individually, clusters the spikes, and then trains a classifier on the clustered data. For each recording channel, the sampled spike waveforms are mapped to their low-dimensional UMAP^17^ embeddings and principal components (PCs). In general, PCA captures global linear variance in the waveform data and can effectively suppress local variability (e.g., high-frequency noise) in its first few dimensions. UMAP, in contrast, preserves local nonlinear structure and manifold geometry that linear methods like PCA inherently miss. By concatenating features from both PCA and UMAP, we construct a feature space that integrates global linear trends with the ability to unfold complex, curved manifold structures. This is one of the notable innovations of our approach, which avoids PCA’s implicit assumption that clusters are roughly Gaussian (ellipsoidal) in PC space and relaxes those shape constraints by incorporating UMAP. This hybrid feature space improves clustering of arbitrarily shaped or overlapping spike clusters and separating morphologically similar or temporally drifting waveforms (**Figure S3**).

After constructing this hybrid low-dimensional feature space of the waveforms, we use an iterative density-based clustering (DBSCAN) approach to detect clusters. After each round of DBSCAN,,, another iteration will be performed on that cluster (**Figure 2A**), if the unit is not refractory or ISIs violation (shorter interspike-intervals than refractory period) is not higher than a threshold (see **Methods**). Note that for each clustering round, the parameters are estimated from its input. We estimate the DBSCAN radius ε by computing each point’s k-th nearest-neighbor distance and selecting the knee of the resulting k-distance distribution (see **Methods**). After clusters are detected, their labels are used to train classifiers which will be used for the data sorting module. We considered three different classifiers including support vector machine (SVM), multilayer perceptron (MLP), and convolutional neural networks (CNN), and evaluated their performance against each other. For SVM and MLP, the classifiers were trained on the first 30 PCs of the waveforms, but for the CNN the raw waveforms containing the spatio-temporal structure of the spike waveforms were used. On average, SVM (with a polynomial kernel, see **Methods** for details of classifiers) outperformed other classifiers. SVM showed ~10x faster training time (**Figure 2B**, top), and significantly higher accuracy (Wilcoxon test, P<0.001) compared with both CNN and MLP when trained on clustered data from real recordings (**Figure 2B**, bottom). This performance was consistent for different sample sizes and test/train splits.

**Figure 2.**
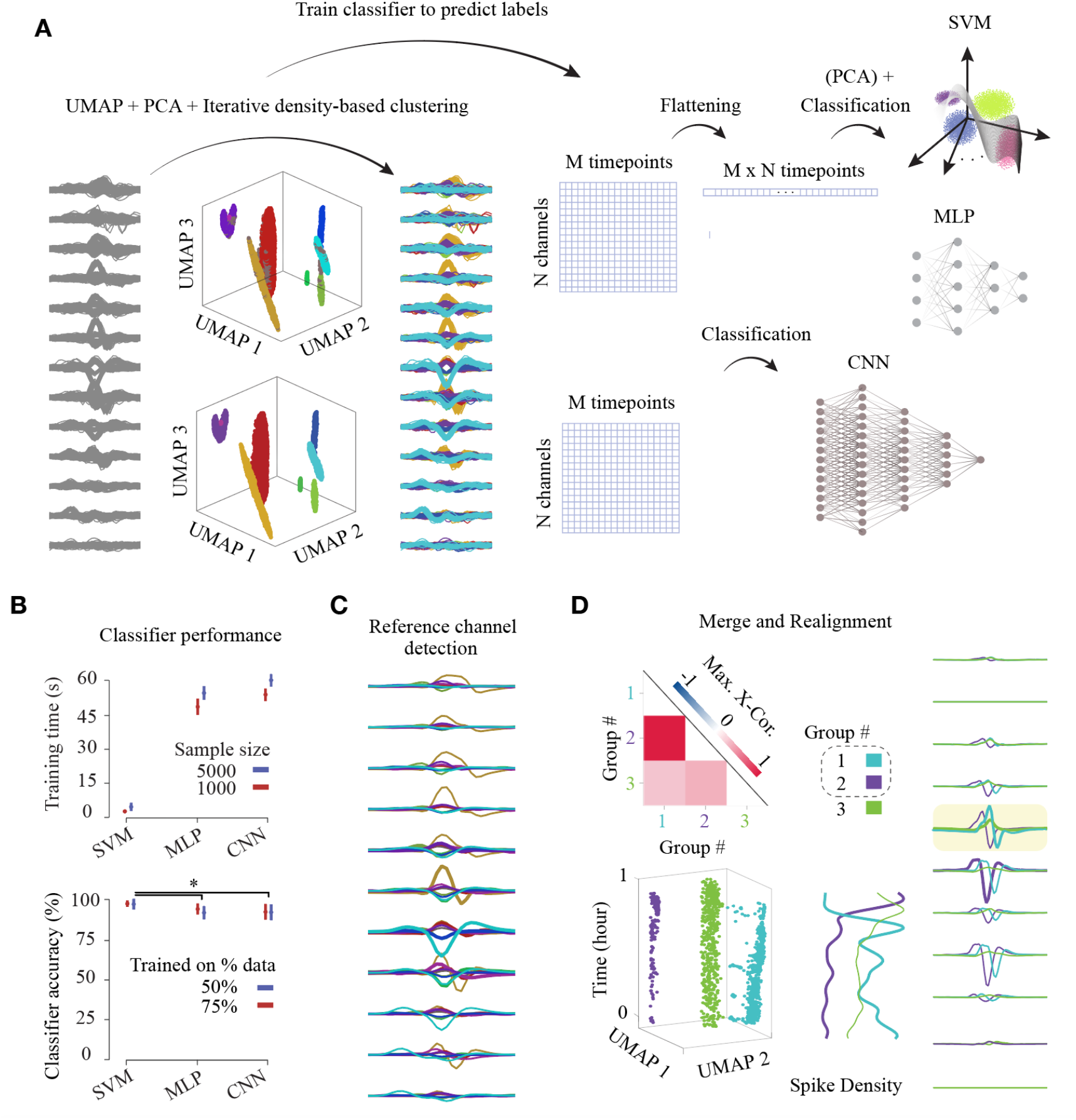
Per-channel spike clustering and classifier training. **A**. Workflow for each recording channel. Extracted spike waveforms are dimensionality reduced using UMAP and PCA, then concatenated into a single feature space for iterative density-based clustering. Clusters that violate the refractory-period or ISI thresholds are recursively split, and low-confidence spikes (gray) are excluded. Final cluster assignments and waveforms train classifiers. The algorithm default classifier is SVM, three classifier types were evaluated: SVM and MLP (trained on principal components), and CNN (trained on raw waveforms). **B**. Classifier benchmarking. Top: training time for SVM, MLP, and CNN on 1,000 (red) versus 5,000 (blue) training sample sizes. Bottom: classification accuracy for two train/test splits (50% vs. 75%). SVMs achieve the fastest training and highest accuracy. Error bars indicate standard error of the mean and significant differences (p < 0.05) are denoted by asterisks. **C**. After clustering a reference-channel is detected for each cluster. Average waveforms are shown for each cluster on neighboring channels. The channel exhibiting the largest absolute peak amplitude is assigned as the reference-channel (thick lines). **D**. An example of cluster merging and realignment. Top left: heatmap of maximum cross-correlations between cluster templates for groups 1–3 flag groups 2 and 3 for merging. Bottom left: 3D UMAP embedding over time showing temporal drift and anti-correlated spike-density profiles of groups 1 and 2. Right: average waveforms for the three groups. Groups 1 and 2 are merged and assigned to the 5^th^ (yellow shaded) channel.

After clustering, to avoid oversplit clusters of spike waveforms (where a single unit with high waveform variability is erroneously divided into multiple clusters), we use a two-tier merging strategy. In the first, *stringent merge*, we merge clusters showing low amplitude variation (less than 5% of peak-to-peak between the mean waveforms) and high zero-lag cross-correlation between their waveforms (r > 0.9). In the second, *looser merge*, we merge clusters with looser criteria on their amplitude variation (15%) if their cross-correlation peaks at nonzero lag (to account for temporal misalignment of multipolar spike waveforms) and if clusters show high negative spike-density correlation (r < −0.8) to capture splits caused by more pronounced spatial drifts (see **Methods** for more details). An example of merge between two misaligned clusters is shown in **Figure 2D**. The two classes show drift (left bottom) and negatively correlated spike density (middle), while showing a high cross correlation value (top left), all of which decides a merge between the two clusters.

For each cluster at any given channel, we define a reference-channel as the channel that shows the largest amplitude for that cluster (**Figure 2C**, reference-channels are shown in thicker lines). This reference channel indicates the channel that each cluster detected at any given channel belongs to. After all channels are processed, the algorithm unifies labels of spikes across all channels, ensuring each cluster has only one reference channel that is the same across all channels containing that cluster. It also assigns cardinal unified labels to all unique clusters detected across the sample data.

### Per-channel classifier-based spike sorting of the data

After processing all sample data, clustering spikes, and training per-channel classifiers, the algorithm executes its third module to sort the full dataset. The full spike sorting is performed in batches. First, each batch is denoised using the same method described earlier. Then, the spike threshold for each channel from the sample extraction phase is applied and spikes are detected on each channel. For each spike detected on a given channel, multichannel waveforms are extracted. Overlapping spikes are passed through a length-adaptive sigmoid suppression function which suppresses all too close spikes on each channel. After extracting the waveforms, if a waveform exhibits its maximum amplitude on the same channel where it was detected, the spike is retained for classification (**Figure 3A**); otherwise, it is temporarily stored for later reference. Retained spikes are classified using the classifier trained on that channel. After the labels are predicted, if the reference channel for the predicted class matches the current channel, the label is accepted, and all associated spike information is stored (including the relabeled and unified label of the spike, channel number, and its index). If the reference channel differs from the detected channel, the algorithm searches for the lookup channel that is associated with the class to determine where the spike belongs, retrieves the corresponding temporarily stored spike at that channel, and reclassifies it in a second pass (**Figure 3A**).

**Figure 3.**
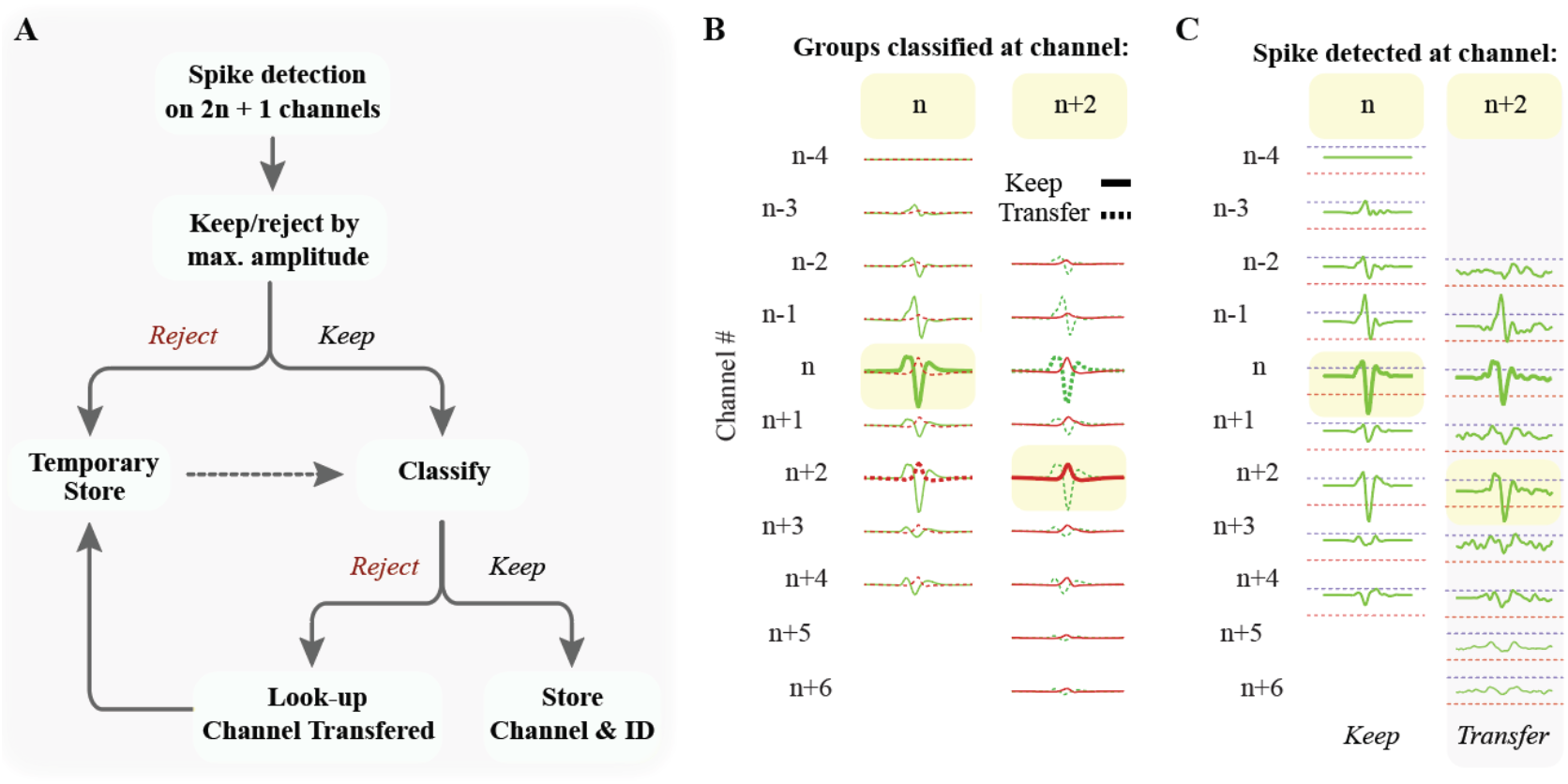
Per-channel classifier–based spike sorting workflow. **A**. Flowchart of the two-pass, per-channel sorting procedure. Spikes detected on each channel (2n + 1 window) are first retained if their peak amplitude occurs on the detection channel ‘n’ or else sent to a temporary store. Retained spikes enter the channel-specific classifier: if the predicted unit’s reference channel matches the detection channel, the unit’s label is stored; if not, the lookup table directs retrieval of the corresponding stored spike for reclassification on its true channel. **B**. Illustration of class assignments for classifiers trained on two nearby channels (n and n + 2). Waveforms whose maximum lies on the local channel are kept (solid traces), while those referencing the other channel are transferred to the channel at which it has its maximum (dashed). Here, the green class references channel n and the red class references n + 2. **C**. Two example spikes from the same neuron showing maximal amplitude on different channels. The left spike peaks on channel n and is correctly labeled in the first pass; the right spike peaks on channel n + 2, is initially classified there, but is redirected back to channel n and in the second pass is correctly transferred to and labeled at channel n.

An example of the same class identified by classifiers trained on two nearby channels is shown in **Figure 3B** depicting retained channels for each class. In this example, the red class has a reference channel at *n+2*, and the green class has a reference at *n*. This keep/transfer process continuously tracks neurons and enables a per-neuron tracking system that, even if a neuron shows drift and its maximum peak is misallocated from its reference-channel, after the first round of classification it will be directed to its reference channel for the second round of classification (**Figure 3C** shows two examples for each condition). After all channels are processed for each batch, the results are unified and written to local or external storage. All files are saved in the HDF5 (Hierarchical Data Format version 5) format.

### Biophysical simulation, drift correction and benchmarking

We examined KIASORT performance on both real and simulated datasets. For benchmarking and comparing its performance against the state-of-the-art method, Kilosort4 ^12^, we used simulated multichannel data. Unlike methods using whole-probe drift tracking ^3^, Kilosort4 implements depth-dependent drift-estimation in which the probe depth is divided into overlapping bands, for each of which a drift trace is found, and data alignment is performed on demand through Gaussian kriging interpolation ^12^. While this band-wise approach relaxes the assumptions about whole probe rigid drifts, it still treats motion as a one-dimensional displacement along the probe axis for each band, which often encompasses multiple neurons. Consequently, it does not correct waveform distortions arising from neuronal morphology (e.g., dendritic or AIS current contributions) or from residual horizontal and oblique probe offsets, especially when the heterogeneity between these neural features as well as the proximity of the probe to the recording units increases.

To better illustrate the limits of such linear-superposition assumptions on drift correction, we built an in silico four-neuron model in which the probe undergoes a 50 µm vertical and a subtle 0.5 µm horizontal displacement (the horizontal shift is 1 % of the vertical) and is then perfectly realigned, “corrected”, along the depth-axis (**Methods, Figure 4A**). We modeled the neurons using biophysically realistic multi-compartmental models of pyramidal cells and interneurons with different morphologies and kinetics (see **Methods**). After drift, channel realignment restores each neuron’s depth position to its baseline by shifting the post-drift waveforms to correct for the vertical displacement (**Figure 4B**). For each before-drift and after-drift-correction condition, we generated a Gaussian cloud of waveforms to compare their low dimensional structures. Our results show that by incorporating even as small as < 1 µm horizontal displacement, the estimated waveforms can differ substantially, showing notable residuals and distinct low dimensional structures, particularly for neurons with faster action potential kinetics (**Figure 4B;** Neuron C). The distortion after drift correction was even worse when the horizontal displacement becomes more notable (> 1 µm), which exacerbated residual distortions and led to more distinct low dimensional structures for all neurons (**Figure S4**).

**Figure 4.**
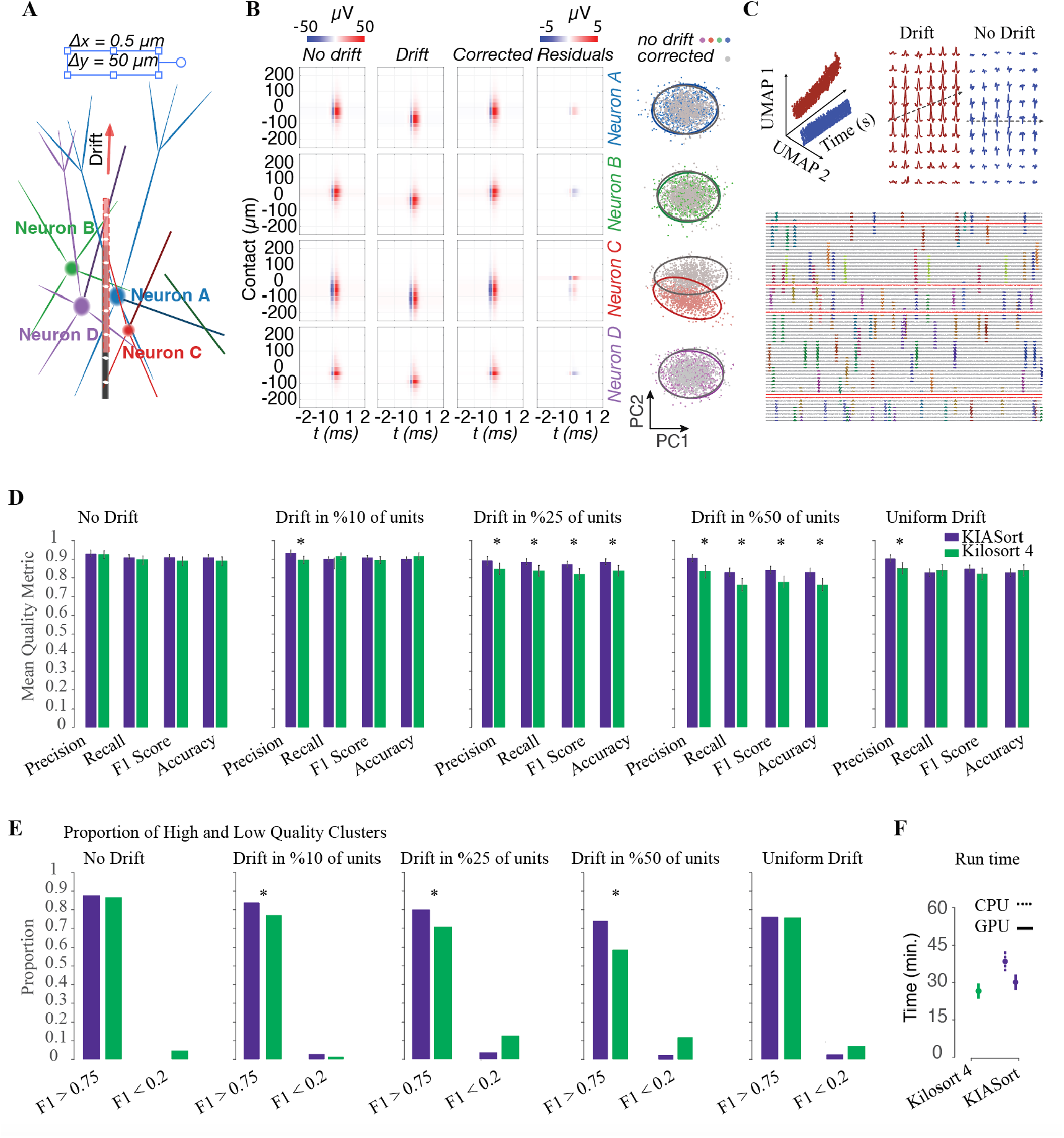
Benchmarking KIASORT performance on simulated datasets with varying drift. **A**. In silico simulation of laminar recording from multicompartmental neuron models, containing pyramidal cells (neurons A and D) and interneurons (neurons B and C). **B**. Extracellular field for spikes of each neuron are modeled before and after a vertical *Δy =* 50µm drift and a small horizontal *Δx =* 0.5µm displacement. Panels from left to right show each neurons’ field before drift, after drift, after drift correction, and the residual from subtracting drift corrected waveform from the waveform before the drift. Rightmost column shows the PCs of waveforms generated for each neuron after drift correction (gray) and before (colored). **C**. Bottom: multi-channel dataset with spike waveforms with variable shapes (uni-, bi-, and tri-phasic) associated with morphology of cell types and, and with different polarities (positive and negative). Sorted spikes are color-coded, and poor-quality channels detect by the algorithm are highlighted in red. An example illustrating two neurons with (brown) and without (blue) temporal drift. Top-left: UMAP embeddings show the temporal trajectories of spike features. Top-right: Waveform changes over time for the drift versus stable unit. **D**. Comparison of mean quality metrics (Precision, Recall, F1 Score, Accuracy) between KIASORT (purple) and KiloSort4 (green) at increasing levels of simulated drift. **E**. Proportion of high-quality (F1 > 0.75) and low-quality (F1 < 0.2) units detected by each algorithm. KIASORT consistently identifies a higher proportion of high-quality clusters across drift conditions. **F**. Average run time for each algorithm when using only CPU (dashed line) or combined with GPU acceleration (solid line). Kilosort4 failed running using only CPU. Error bars indicate standard error of the mean and significant differences (p < 0.05) are denoted by asterisks.

For our benchmarking, therefore, we implemented per-neuron drift by modulating spike amplitude along the depth axis (see **Methods**). First, we generated datasets using biophysically realistic spike waveforms containing both negative and positive spikes with uni-, bi-, and tri-phasic waveforms representing morphology variabilities and different electrode proximities (see **Methods**). We then added background noise by generating a signal whose frequency spectrum matches that of Neuropixels recordings in mouse cortex, with some channels modeled to mimic the spectral content of silent or poor-quality contacts (**Figure 4C**). Each neuron could drift up to 50 µm from its original location. Drift was implemented either as continuous Gaussian waveform changes, stepwise neuron displacement, or continuous displacement with nonlinear distortions (see **Methods**). An illustrative pair of UMAP trajectories depicts one neuron that drifts and another that remains stationary, highlighting the temporal evolution of waveform space (**Figure 4C**).

We evaluated both KIASORT (purple) and Kilosort4 (green) on simulated datasets with varying proportions of drifting units, uniformly distributed along the depth axis. For Kilosort4, we optimized the detection threshold and parameters to maximize performance (in general, lowering the threshold tended to introduce false positives). For each detected unit, we computed precision, recall, accuracy, and F1-score (see **Methods**). Our simulation results showed KIASORT performing comparably to Kilosort4 on no-drift and uniform-drift conditions, while outperforming Kilosort4 in per-neuron drift conditions, with the performance gap widening as drift variability between units became more pronounced (**Figure 4D**). These performance differences were consistently evident when waveform changes were nonlinear, or drift was implemented nonlinearly (**Figure S5**). Consistent with this finding, the proportion of high-quality units (F1-score > 0.75) detected by each algorithm showed KIASORT identified a significantly higher number of high-quality units, particularly under conditions with greater variability in drift across units (**Figure 4E**). Overall, while Kilosort4 shows decent performance on conditions with no drift or when neurons within similar localities show consistent drift, our geometry-free and per-neuron tracking shows significantly improved performance, particularly when these assumptions do not hold true in real-world complex neural recordings. We also tested the run time for both KIASORT and Kilosort4 on 1 hour of 384-channel data. KIASORT showed comparable performance to Kilosort4 when using GPU. On the other hand, while we were unable to run Kilosort4 using only CPU, KIASORT still showed real-time performance with CPU alone (**Figure 4F**).

### KIASORT performance on mouse and monkey high-density neural recordings

We ran KIASORT on datasets recorded from different probes. We used data from the caudate nucleus, anterior cingulate cortex, and lateral prefrontal cortex in the monkey using a dual-shank diagnostic biochips (DBC) probe, and datasets from the Allen Institute free-access online repository, recorded from mouse cortex. In all datasets, we found that our noise removal method could eliminate any shared noise or signal components evident on a high number of channels, even when the noise level was prominent (**Figure 5A**). Our algorithm yielded ~2x spike/channel, showing ability to detect even high-firing fast-spiking neurons (fr > 40 spike/sec) that often do not show dips in their spike train correlograms. It also was able to track neurons and detect units with different waveform shapes (see **Figure 5B** for an example). Similar performance was observed for recordings in mouse (**Figure 5C**). In all datasets, we also documented per-neuron drift as a likely event in neurophysiological recordings. As illustrated in **Figure 5D** as an example, from three highly isolated nearby neurons, only one showed drift with a pattern different from the other neurons. With its geometry-free design and per-neuron tracking sorting system, KIASORT successfully tracked and identified these neurons independently in tested real datasets in recordings from different brain areas, including Striatum, lateral prefrontal cortex, and anterior cingulate cortex which contains neurons with different morphologies (**Figure S6**). In the example DBC and Neuropixels recording, KIASORT detected 236 (from 128 channels) and 733 (from 384 channels) and for active channels showed on average 1-2 high quality isolated single units. In addition, while a direct comparison between KIASORT and Kilosort4 on real data without ground truth is not a valid benchmark, we compared the sorting performance between the two methods. In a recording session from the primate striatum, we computed the spike amplitude on the main channel for both Kilosort4 and KIASORT, and defined SNR consistently across methods. We then quantified the fraction of refractory units (defined as those with an inter-spike interval (ISI) violation rate below 0.2%) for each method. We observed that KIASORT detected around 5% higher proportion of refractory units among those with high signal-to-noise ratio (SNR) than Kilosort4 (**Figure 5E**).

**Figure 5.**
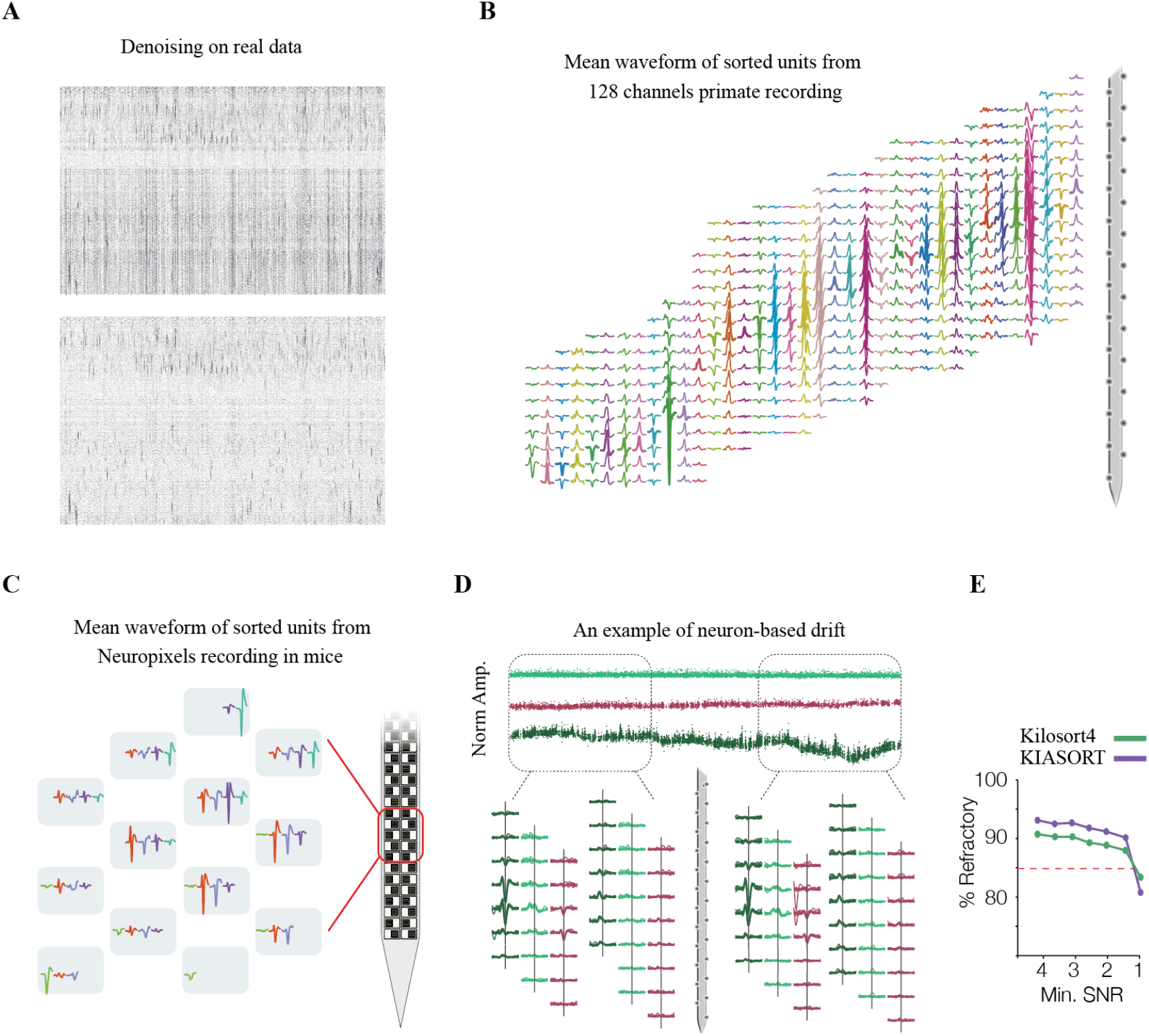
KIASORT results on real data. **A**. An example of two-step denoising applied to raw primate probe recordings, showing the raw voltage trace with variable artifacts on different channels (top) and the cleaned signal after denoising (bottom). **B**. Example mean multichannel spike waveforms of units sorted from a 128-channel, dual-shank DBC probe in primate striatum. **C**. Example mean multichannel spike waveforms of units sorted from a Neuropixels probe recording in mouse cortex (Allen Institute dataset). **D**. An example of neuron-based drift in a primate DBC probe: one unit (dark green) exhibits progressive spatial drift over time, while two other units (light green and red) remain stable throughout the recording. **E**. Fraction of refractory units within different minimum SNR value compared between KIASORT and Kilosort4 for the same session as in **D**.

## Discussion

We introduced KIASORT, a novel spike-sorting algorithm for high-channel recordings, and showed that it addresses key limitations of current methods. KIASORT implements a per-neuron tracking system and uses a geometry-free approach to effectively handle neuron-specific drift and nonlinear waveform changes that challenge existing spike-sorting algorithms. Our results demonstrate that KIASORT outperforms the current state-of-the-art method, Kilosort4, particularly under conditions with pronounced per-neuron drift variability (**Figures 4D–E**), which is common in real neurophysiological recordings (**Figure 5D**).

One of the key innovations of KIASORT, compared with existing methods, is its hybrid use of nonlinear UMAP combined with linear PCA for dimensionality reduction (**Figure 2A**). While previous approaches rely primarily on linear embeddings ^6,7^ with a priori assumptions about the geometry of the cluster feature space, integrating UMAP’s nonlinear manifold preservation makes KIASORT better suited to arbitrarily shaped clusters and temporally changing waveforms. This hybrid approach can also handle complex drift manifolds that might otherwise be oversplit or merged inappropriately ^10,16^.

Another innovation and unique feature of KIASORT is its relaxation of probe-geometry assumptions in favor of a per-neuron, data-driven tracking approach that follows each neuron independently. Compared to earlier correction methods ^3^, more recent approaches (from Kilosort2 onward) have implemented piecewise drift-tracking and correction models ^12^; however, these methods still assume that neurons in close spatial proximity experience similar drift patterns and thus fail to account for drift arising from horizontal probe displacement, oblique insertion, or heterogeneous neural orientations. Our biophysical simulations demonstrate that even small deviations from one-dimensional motion can substantially distort waveforms in ways that cannot be corrected by even perfect one-dimensional displacement models (**Figure 4B**). KIASORT’s hybrid approach for clustering, channel-specific classifier training, and neuron-based tracking system allow it to adapt to these complex patterns without imposing geometric constraints. The performance advantage is particularly pronounced when the variability of nonlinear waveforms or the heterogeneity of drift increase in the neural population (**Figure 4D–E**), with KIASORT identifying 5–15% more high-quality units than Kilosort4.

The modular structure of KIASORT (**Figure 1**) is another advantage, particularly for tracking neurons that remain consistently available across multiple recording sessions. KIASORT separation of the sorting process into three distinct modules, allows users to sample data and train classifiers on one dataset, then apply them to others by running only the third module (sorting the data). This modular design is integrated into a user-friendly GUI with editing features that facilitate data inspection, spike sorting, and post-hoc curation within a unified framework. This all-in-one platform is another development of KIASORT that is currently missing from existing spike-sorting pipelines ^18,19^.

Despite these advances, it is worth noting that, like any algorithm, KIASORT faces challenges and limitations that we outline here. A key challenge occurs when drift is so severe that a neuron completely vanishes and becomes indistinguishable from baseline noise, so that no spikes are detected on the reference channel. In such cases, the algorithm may split a single neuron into multiple units, which then might require post-hoc manual curation and merging which can be done within the same sorting platform (**Figure S1**). Another limitation is when neurons have very low firing rates (< 0.1 Hz), which might be missed entirely during the sampling phase and thus not identified in the clustering stage. Additionally, severe but transient noise conditions, such as electrical stimulation artifacts that saturate the amplifier and cannot be effectively removed, do not necessarily prevent spike detection but, if they occur frequently, can result in false-positive unit detections, which is addressable with post-hoc manual curation.

In summary, KIASORT presents a novel and fundamentally restructured spike sorting algorithm, showing significant advances in spike sorting for multichannel recordings, particularly for recordings with complex drift patterns. Its geometry-free design and per-neuron tracking approach make KIASORT less dependent on prior assumptions, making it suitable for non-linear variabilities in waveform shapes, and different probe design.

## Materials and Methods

### Denoising

To remove any shared noise across a large number of recording channels, we applied a percentile-template regression that preserves neural spikes waveforms intact. The denoising can be GPU-accelerated when supported. For each raw signal X ∈ *R*^*N*×*M*^, with N channels and M time samples, we first compute the percentile at each time point:

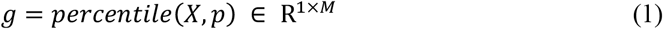

Where *p* is the percentile value used to form the noise template. By default, it is set to 50 (meaning 50% of channels need to show the noise pattern) but can be adjusted by user if the noise pattern is different in their datasets (e.g., when noise is present on less than half of channels). Then each template is moved to the center by:

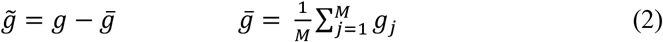

and each channel data centered by:

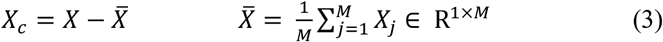

then the correlation between each channel and the noise template as:

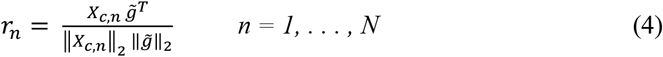

for each channel *n*, the algorithm finds a regression gain as:

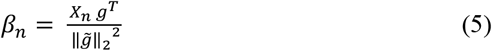

and an amplitude gain as:

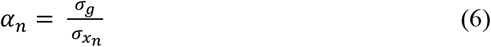

where σ_g_ is the standard deviation of the noise template, and 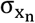 is the standard deviation of channel n. For all channels exceeding both an amplitude gain threshold and a correlation threshold, the algorithm subtracts the noise from the signal by:

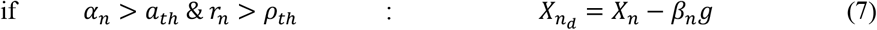

where *ρ*_*th*_ is the correlation threshold, by default set to 0.4, and *α*_th_ is the amplitude gain threshold indicating a fraction of root-mean-square (RMS) amplitude, by default set to 0.05 to conservatively ignore normal spiking activity ^3,20^.

### Signal Quality Check

To assess channel quality, we compared the power in the spike band (1.5 kHz < f < 4.5 kHz) with the power in the low-frequency band (0.5 kHz < f < 1.5 kHz) and the high-frequency band (4.5 kHz < f < f_s_/2). We then normalized the power in each band by its bandwidth and computed a ratio that indicates whether a channel is dominated by low-frequency or high-frequency noise.

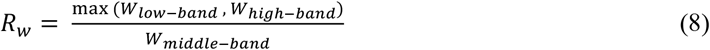

The ratio is then transformed to scale to the bands by:

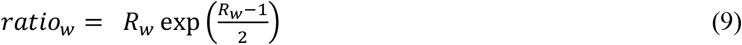

If the ratio is too high, it indicates that the signal is dominated by either low- or high-frequency noise without prominent mid-band power ^22,24^. We set the ratio threshold to the smaller of a fixed value (3 by default) and the mean ratio across all channels plus three standard deviations. We evaluated this approach on simulated multichannel datasets as described above and on real recordings from non-human primates (**Figure S2**). For the spike sorting, any channel not passing this quality check will be excluded from spike detection and waveform extraction.

### Sample extraction

For the sample-extraction phase, the algorithm selects chunks of t-second duration signals (by default 1 second) for all channels, *X*(*t*_*i*_: *t*_*i*_ *+ t*). A total of M sample chunks (by default 180) is extracted from regularly spaced segments of the data. Users can change these values depending on the data and also set a higher limit on the maximum time point up to which samples will be extracted. Once all sample chunks have been extracted, they are concatenated into signal *X*_*sample*_ ∈ *R*^*N*×*Mt*^ which is then used for subsequent sample spike detection.

### Spike detection

For each recording channel, we detect all spikes that exceed a threshold set by multiplying a scale factor by the median absolute deviation (MAD) of that channel’s signal. The algorithm first bandpass filters the data between 300 Hz and 8000 Hz (adjustable for recordings with lower sampling rates), then calculates the MAD threshold as follows:

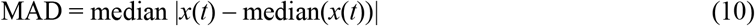

The threshold is by default set to 7 × MAD, which works well for many datasets. However, users can adjust this scale factor in the GUI or by scripting depending on their data. In general, MAD thresholding is robust to outliers and noise. However, to prevent threshold inflation on channels with very high spiking activity, the algorithm first detects spikes using MAD thresholding, excludes those spikes, and then reapplies MAD thresholding to compute the final threshold. Spikes exceeding this threshold are detected. For detected spikes that occur too closely in time, only the one with the highest peak amplitude is retained and other will be disregarded. The minimum interval between spikes is set to 0.375 ms by default, but if the distribution of inter-spike intervals (ISIs) shows its largest peak between 0.375 ms and 0.75 ms, the interval is adaptively increased to match that peak and preventing double-counting of multiphasic spikes with widely but regularly spaced peaks ^6^.

### Waveform extraction

For each detected spike on a given channel, we define an extraction window of fixed length (default 1 ms). This window is centered on the spike’s most extreme positive or negative peak, and the waveform within this window is extracted across the specified neighboring channels. By default, KIASORT extracts waveforms from 15 channels (seven above and seven below the center channel), which for a Neuropixels probe with 20 µm site spacing spans ~280 µm. Users can adjust the number of channels to fit their probe (a range covering 200–400 µm is recommended). During the sample-extraction phase, all extracted waveforms are saved for every channel. However, in the main sorting phase, only waveforms for spikes assigned to the center channel are saved (optional if selected by users).

### Handling overlapping spikes

When two spikes occur very close in time (i.e., when the difference between consecutive main spike indices is less than L), the extracted windows overlap. During the sample-extraction phase, we consider only spikes that have no overlapping spikes detected within their waveform window on any neighboring channels. This prevents splitting a single neuron into multiple clusters due to consistent overlaps with other neurons during clustering.

For the main data-sorting step, however, to compensate for overlapping spikes, the algorithm first computes the overlap length based on the inter-spike interval and then applies a smoothly varying sigmoid window to attenuate the contribution of the overlapping regions from neighboring spikes. If the time difference between consecutive spikes is Δt (in samples) and Δt < L, the overlap length is computed as:

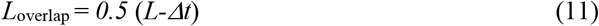

For each possible overlap length *i*, (with 1 ≤ *i* ≤ *L*), we find a sigmoid window by:

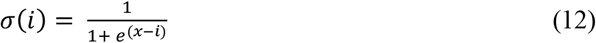

where x is sample point form spike center. Two modified windows are then defined: the left-side window equals *σ(i)* over its left half and 1 over its right half, while the right-side window equals *σ(i)* over its right half and 1 over its left half. Depending on which side the overlap occurs, the corresponding left- or right-side window is multiplied elementwise with the waveform.

### Sorting Samples

In this step, the algorithm loads the sample data previously extracted and saved for each channel. Each channel is then processed individually, spike waveforms are preprocessed, and features are extracted. Spikes are clustered, and classifiers are trained on the clustered data. After processing each channel, the algorithm unifies clusters across channels and assigns a reference channel to each cluster.

### Waveform preprocessing and feature extraction

For each channel, spike waveforms were extracted as T time samples centered on detected threshold crossings across C channels. For each detected spike i, the extracted waveform is a matrix *W*_*i*_ ∈ *R*^*C×T*^ spanning *±τ* ms around the spike peak, where *τ* is half of the clustering spike duration. To prepare for dimensionality reduction, each waveform matrix is first flattened into a one-dimensional vector *vec(W*_*i*_) ∈ *R*^*1×C*.*T*^ and multiplied by a Hamming window function to weight stronger on the central portion of the spike:

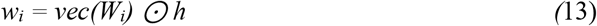

where h ∈ R^*1×*C·T^ is the Hamming window and ⊙ denotes elementwise multiplication.

We used a hybrid approach combining linear and nonlinear dimensionality reduction to form a feature space that captures both global and local structures in the spike waveforms. First, the algorithm applies Principal Component Analysis (PCA) to the preprocessed waveforms:

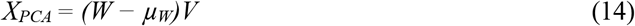

where W ∈ *R*^*N*×(*C*·*T*)^ is the matrix of *N (number of spikes)* flattened spike waveforms, *µ* is the mean waveform vector, and V ∈ *R*^(*C*·*T*)×*n*^ is the first *n* PCs.

For the non-linear part, the algorithm uses Uniform Manifold Approximation and Projection (UMAP)^17^ as:

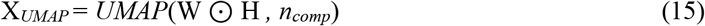

where *H*∈ *R*^*1×C*.*T*^ is the Hamming window applied to all waveforms. *H* is formed by concatenating C individual Hamming windows, each of length T sample points, and *n*_*comp*_ is the number of UMAP components. While PCA largely represents the global variance across the entire waveform, UMAP is more weighted by the central portion of each spike to contain spatial information. It preserves both local and global topological structures and the relative distances between waveforms by projecting them into a low-dimensional nonlinear manifold.

The final feature space used for clustering is constructed by concatenating the normalized UMAP components, the first three PCA components, and the peak amplitude values, as:

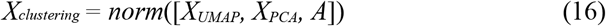

where each feature dimension is min-max normalized to the range [0,1].

### Density-based clustering

The normalized feature matrix X_*clustering*_ ∈ R^*N*×*D*^, is then used for clustering using an iterative DBSCAN method^21^. For each sample *x*_*i*_ ∈ R^*D*^, DBSCAN defines a neighborhood as

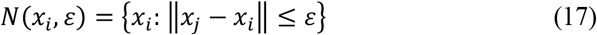

and a point is considered a core point if

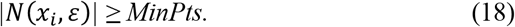

where *MinPts* is the minimum-points parameter in DBSCAN, set according to a minimum firing rate of 0.1 spikes/sec, and bounded between 0.1 % (lower bound) and 1 % (upper bound) of the total data points.

For clusters with ISI-violation ratios exceeding a threshold (indicating potential mixing of multiple units), we recursively apply DBSCAN with adjusted parameters until either the clusters satisfy the refractory-period constraint or a maximum recursion depth of four is reached. After all recursions, we unify the labels, and any noisy clusters or points identified as noise are assigned a “–1” label.

### Parameter estimation for DBSCAN

To optimize DBSCAN parameters, we use a knee detection method^23^ to estimate optimal epsilon. We use k as the number of datapoints, and then compute the *k*-distance for each point x_*i*_ as:

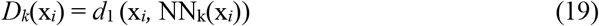

where *NN*_*k*_*(x*_*i*_) is the *k*th nearest neighbor using Manhattan distance. After sorting distances, we detect the “knee” in the sorted distance curve, *D*_(1)_ ≤ *D*_(2)_ ≤ ⋯ ≤ *D*_(*n*)_, by maximizing the perpendicular distance to the line connecting the first and last points as:

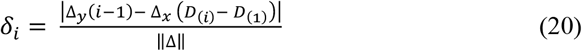

where Δ_*x*_ = *n* − 1, Δ_*y*_ = *D*_(*n*)_ − *D*_(1)_, and **Δ** = (*n* − 1, *D*_(*n*)_ − *D*_(1)_). The epsilon is then set as:

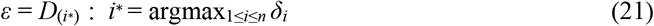

### Cluster merge

When the clusters are labeled for each channel, the algorithm post-process these clusters to avoid overspill. For clusters with a mean waveform *w*_*s*_ ∈ *R*^*M×T*^ and spike density of time series ds, initial labels *l*_*0*_, and sample count ns, the algorithm performs the following steps:

First, for each pair of clusters *(i,j)*, it computes the cross-correlation between their mean waveforms and identifies the maximum correlation coefficient:

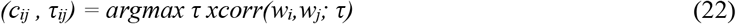

Where *c*_*ij*_ is the maximum correlation coefficient peak, and *τ*_*ij*_ is the time lag at which the peak value occurred.

It computes the amplitude variability by:

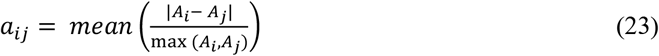

where *A*_*i*_ is the maximum absolute amplitude of *w*_*i*_. Then it computes the correlation of spike density by:

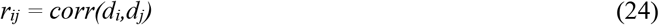

After computing these metrics, clusters can be merged using two strategies: *stringent merge* and *loose merge*. In the *stringent merge*, clusters are merged if they show low amplitude variation, and high zero-lag cross-correlation between their mean waveforms:

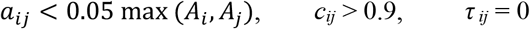

For the *loose merge*, clusters are merged if they satisfy:

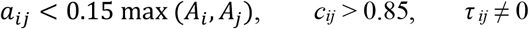

where the nonzero lag criterion accounts for temporal misalignment of multiphasic spike waveforms.

Lastly, the algorithm checks for *drift-based merge* where clusters show:

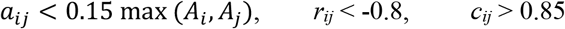

to merge split clusters caused by pronounced spatial drift.

For all merge operations, the resulting cluster’s ISI-violation rate (i.e. the percent of interspike intervals within the refractory period) must not increase by more than 0.1 % ^24,25^.

### Unifying clusters across channels

After clustering spikes on all channels, we assign a reference channel to each cluster. The reference channel is the channel on which the cluster’s mean waveform shows its highest absolute peak and determines which classifier will be used during the sorting stage. For each cluster on a given channel, the reference channel can be either the original channel (in which case the cluster is retained) or a different channel (in which case it is transferred). When a cluster’s reference channel differs from its original channel, the algorithm checks all channels detecting that cluster and enforces consistent reference-channel assignments. If the same cluster is assigned reference channels on two or more channels (defined by more than 50% overlap in spike indices), the channels are unified and only the one with the highest peak value is retained. Finally, if a cluster’s reference channel is not its original channel, the algorithm iteratively confirms that the cluster exists on the reference channel; if not, it relocates the cluster to the ultimately retained reference channel.

### Training classifier

Following unsupervised clustering, we train a supervised classifier for each channel to classify new spikes during the sorting stage. We evaluated three classifier types including support vector machine (SVM)^26^, multilayer perceptron (MLP), and convolutional neural network (CNN)^27^, to compare their performance (**Figure 2A**). KIASORT supports all three, but the default is SVM due to its computational efficiency, higher accuracy, and roughly tenfold faster runtime (**Figure 2B**).

For training, we use either the raw waveforms (for CNN) or the PCA-reduced feature vectors (for SVM and MLP):

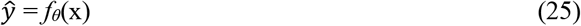

where x is the PCA scores of the flattened waveforms or the raw waveforms, and *f*_*θ*_ denotes the classifier with parameters *θ*. During actual sorting, each classifier is applied to all sampled spikes; however, for benchmarking, we trained and tested using 50/50 and 75/25 train/test splits.

SVM: We use a degree-3 polynomial kernel (radial-basis-function kernels are also supported, but the polynomial kernel performed best). To exclude potential outliers, the SVM leaves 10 % of the sampled spikes out as noise during training. All hyperparameters were tuned for optimal accuracy.

MLP: The MLP has an input layer, four hidden layers with 2048, 1 024, 128, and 64 units, and an output layer. It uses mean squared error loss, a dropout rate of 0.1, an initial learning rate of 10^−4^, and drops the learning rate by a factor of 0.1 every 10 epochs. It is trained on PCA-reduced waveform features. All hyperparameters were tuned for optimal accuracy.

CNN: The convolutional network comprises four blocks with 16, 32, 64, and 128 filters. Training used a batch size of 128, a kernel size of 3, pooling size of 2, L2 regularization at 10^−4^, and matched the MLP’s dropout and learning-rate schedule. All hyperparameters were tuned for optimal accuracy.

### Sorting Data

After extracting samples, clustering the sampled spikes, training classifiers on each channel’s spikes, and unifying spike labels, the algorithm runs the third module to integrate the acquired knowledge and sort the full dataset (**Figure 3**).

To make the computation more efficient, multi-channel neural recordings are segmented into temporal chunks of duration *T*_chunk_ (1-2 minutes are optimal). To avoid boundary artifacts, each chunk is extended by a small margin *T*_*margin*_ (2 seconds), and any spikes detected within this margin are excluded. Similar to the sample-extraction step, raw data in each chunk are bandpass filtered and denoised using the noise-template regression. For each channel, spikes are detected exactly as in the sampling phase, and waveforms are extracted for each spike. Overlapping spikes are then attenuated using the sigmoid window method described previously.

For each detected spike, the algorithm checks whether it has the maximum peak amplitude on its current channel, if so, it is retained for classification there; otherwise, it is temporarily stored for cross-channel transfer. Each recording channel has a pre-trained classifier *fi* : R^*k*^ → {1,…,*N*_*ch*_} that was trained on PCA-reduced features during the sampling phase. Given a spike’s feature vector x on channel i, the classifier outputs a label 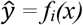 and posterior probabilities *p(y=k* | *x)* to quantify the confidence. Otherwise, the spike is transferred to its reference channel: the algorithm searches among the temporarily stored spikes on the reference channel for a spike whose detection times t’ satisfy |*t* ™ *t*′| *<* Δ*t* (Δ*t* = 0.75 ms), then reclassifies the spike using the reference channel’s classifier. To support this cross-channel lookup without excessive memory use or compute overhead, channels are processed in a sliding-window by loading classifiers for the *2N* channels before and after each channel (where *N* equals the number of channels used for waveform extraction). After all channels have been evaluated for each chunk, the results are stored in an HDF5 file containing spike indices, their labels, the label on each channel, their channel, their peak amplitude on the reference channel, and, optionally, multichannel waveforms.

### The unified GUI platform

KIASORT provides a unified graphical user interface (GUI) that integrates data visualization, inspection, and parameter settings for sorting on its main tab (Figure S1). The GUI includes three additional tabs:

Channels: Enables users to inspect raw data, manually include or exclude channels, and adjust the MAD threshold via a slider.

Curation: Lets users load sorted results and perform post hoc processing; mark unit isolation types, merge or remove channels, realign waveforms, and evaluate waveform similarity, and explore cross-correlograms, presence ratios, ISI distributions, and feature metrics before saving curated outputs (Figure S1).

Clusters: Offers a sanity-check view of clustering results for each channel, allowing users to fine-tune parameters in cases of atypical datasets.

### Biophysical Modeling of Probe Drift Effects on spike waveform

We developed a biophysically plausible model to investigate the effects of laminar probes drift on extracellularly recorded action potentials. We model multicompartmental neurons with proper cable dynamics, realistic membrane biophysics, and spatially extended current sources to capture the complex, nonlinear relationship between probe movement and spike waveforms.

### Multicompartmental neuron models

In the modeling we considered four distinct neuron morphologies, representing major cell types in cortical circuits, each with biophysically plausible but simplified geometry and compartmentalization (**Figure 4A, Table S1**):

Pyramidal Neurons (A, D): Large spherical somata (20–22 µm diameter), apical dendrites extending 180–200 µm at 70–100° angles, basal dendrites 100–120 µm long at random orientations, and axon initial segments 180–200 µm in length.

Stellate Neuron (B): Medium soma (18 µm diameter), radially oriented dendrites 100–130 µm, axon 150 µm in length.

Interneuron (C): Smaller soma (15 µm diameter), shorter dendrites 70–90 µm long, and a thin 1.5 µm axon 120 µm in length.

Each compartment was modeled using standard cable theory ^28^, in which the membrane potential along each compartment obeys the cable equation:

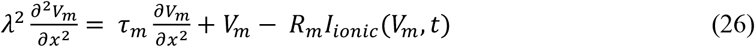

where λ is the space constant, τ_m_ is the membrane time constant (1.0 ms), R_m_ is the specific membrane resistance, and *I*_*ionic*_ denotes the nonlinear ionic currents.

### Membrane ion channel dynamics

The membrane potential dynamics are modeled by nonlinear activation of sodium and potassium channels by adapting biophysically plausible kinetics from the Hodgkin–Huxley equation ^29,30^:

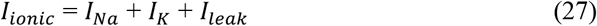

Sodium channel activation followed a sigmoidal relationship with membrane potential:

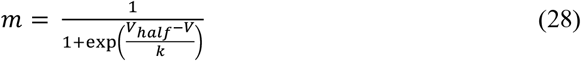

where *V*_*half*_ = −40 *mV* denotes the half-activation voltage and k = 6 indicates the slope factor. Channel saturation was incorporated using a hyperbolic tangent nonlinearity:

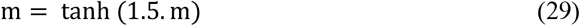

Potassium channel activation showed gating and slower kinetics as:

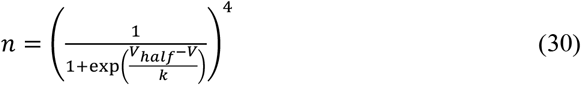

whit *V*_*half*_ = −55 *mV* and k = 8.

The resulting compartment-specific currents were modeled for the soma by:

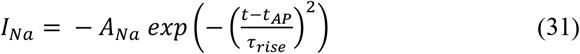

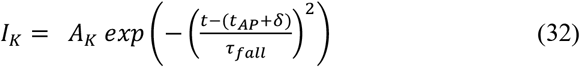

where *t*_*AP*_ is the action-potential timing, τ_rise_ and τ_fall_ are the compartment-specific membrane time constants, and *δ* is the delay in potassium-current activation. We also incorporated cell-type specific differences in the kinetics of these currents with interneurons showing faster kinetics (shorter *τ*_*rise*_ values) than pyramidal cells ^31,32^.

### Field potential calculation

We calculated the extracellular potential generated by each compartment using the line source approximation ^14,28^, which estimates each neuronal segment as a line current source within a homogeneous volume conductor by:

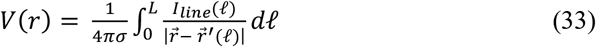

where σ = 0.3 S/m is the extracellular conductivity, *I*_*line*_ is the transmembrane current per unit length, *r* is the electrode position, and *r′(ℓ)* denotes positions along the compartment of length *L*.

For computational efficiency, this integral was solved analytically for a line segment carrying constant current. For a segment running from **r**1 to **r**2 at a perpendicular distance *h* from the electrode, the extracellular potential is:

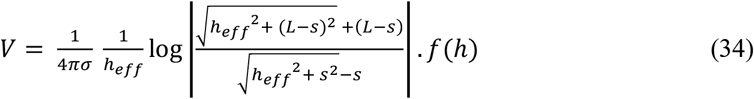

Where 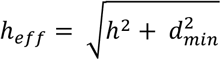, incorporating a minimum effective distance of *d*_*min*_=10 µm to avoid singularities, s is the longitudinal projection of the electrode onto the segment, and

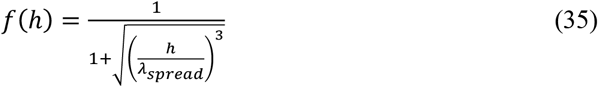

a spatial-decay function with *λ*_*spread*_ = 60 µm.

For very short segments or point-current sources, this simplifies to:

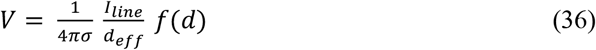

where *d* is the distance from the source to the electrode.

We incorporated a critical biophysical phenomenon, finite propagation velocity of extracellular fields, into our model. Unlike simplified models assuming an instantaneous transmission for neural signals, we account for spatiotemporal signal delays to estimate how electrode drift, combined with propagation time, can distort post-drift spike waveform reconstruction. In brain tissue, electrical signals travel at roughly 0.1–10 m/s depending on myelination, frequency content, and other medium related factors. ^33,34^.

To simulate finite propagation times from neuron to electrode, we consider channel-dependent delays:

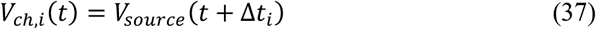

where 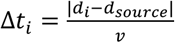 is the propagation delay between the channel *i* and the nearest channel to the source. Here, *v* is the propagation velocity (5 m/s), which leads to delays of about ~1 µs per channel distance for high-frequency components ^34^. This ensures that signals reach different recording sites with realistic temporal offsets.

### Probe drift and waveform drift correction

A probe drift was implemented by relocating the probe on the vertical axis by *Δy* and on horizontal axis for shear effect by *Δx* displacement (Figure 4A). To avoid any inaccuracies for the reconstruction, we shifted the probe only by multiples of channel spacing (25µm). In our simulation, we used *Δy =* 50 µm two different *Δx* values of 0.5 µm, and 5µm which represents an angle of ~0.5^·^ and ~5^·^ respectively.

For each probe, we then correct the drift by simply shifting its channel up or down to match the location of the probe before drift.

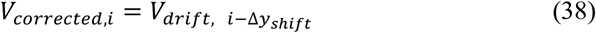

All waveforms for each condition were aligned to their peak-amplitude time (t = 0) for further comparisons.

To examine the drift-correction efficacy in the context of spike-sorting we generated 1000 waveform variants for each neuron before drift and after the drift-correction random amplitude scaling and adding background noise. We then used PCA to reduce waveforms and visualize the separability of clusters in their feature space before and after correction.

**Table S1.** Simulation parameters used for the biophysical modeling of the probe drift.

| Parameter | Value |
| --- | --- |
| Extracellular conductivity ($\sigma$) | 0.3 S/m |
| Probe geometry | 50 channels with 25 $\mu\text{m}$ distance |
| Time vector ($t$) | $[-2, 2]$ ms; 0.005 ms resolution |
| Membrane time constant ($\tau_m$) | $1.0 \times 1.5$ ms |
| Sodium channel time constant ($\tau_{rise}$) | $0.12 \times 1.5$ ms |
| Potassium channel time constant ($\tau_{fall}$) | $0.25 \times 1.5$ ms |
| Minimum effective distance ($d_{min}$) | 10 $\mu\text{m}$ |
| Spatial spread factor ($\lambda_{\text{spread}}$) | 60 $\mu\text{m}$ |

### Simulating Multichannel Extracellular Spike Data

We simulated multichannel extracellular data by combining background noise and spike data. First, we estimated the background signal using spectral features from real Neuropixels recordings. We modulated the FFT of the signal by a coefficient applied to the frequency components, and then incorporating a spatial factor so that nearby channels exhibited greater similarity in their low-frequency variabilities. For each recording channel, LFP noise was modeled as the sum across logarithmically spaced frequency bands. For band b, with frequency range [*f*_1,*b*_,*f*_2,*b*_], we compute:

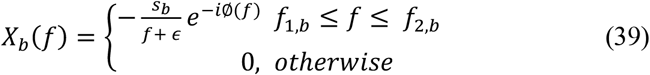

where *S*_*b*_ is the scaling factor, *ϵ* is a small constant to avoid singularity at *f* = 0, and *ϕ*(*f*) is a random phase uniformly distributed over [0,2*π*), but constrained by a spatial factor that causes nearby channels to exhibit greater phase alignment in their low-frequency bands. The time-domain signal for band b is then obtained via the inverse Fourier transform:

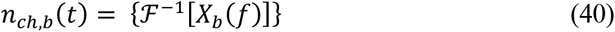

where *n*_*ch,b*_ is the signal for frequency band b, at channel n. The raw background signal is then found through sum over all frequency bands:

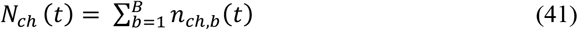

### Spike generation

For each channel we randomly designate 0 to 3 neurons to show their peak amplitude. Each neuron is then assigned a firing rate *r* from a lognormal distribution^35^:

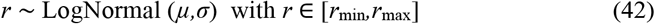

Where *r*_min_ *and r*_max_ min and max firing rate set between 0.25 and 40 spike/sec respectively. Spike times follow a Poisson process with exponentially distributed interspike intervals:

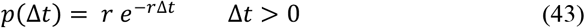

For neurons, a refractory period Δ*t*_ref_ = 2 ms is enforced such that:

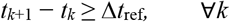

We then modeled spike waveforms as monophasic, biphasic, or triphasic shapes to capture variations in morphology and electrode proximity. Monophasic spikes were modeled as:

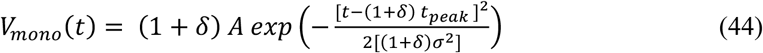

biphasic spikes as:

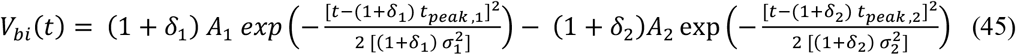

and triphasic spike:

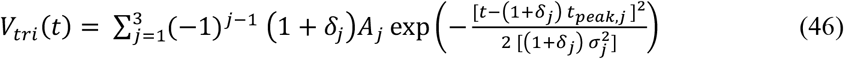

where *δ*_*j*_ is small random perturbation introducing variability, A_j_ is the amplitude for each phase extreme, and t_peak,j_ is time to peak for each extreme. Each spike waveform is further perturbed by multiplicative noise (scaling by 1 + *η* with 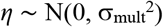)) and an additive noise 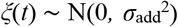):

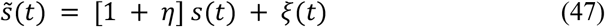

For each spike waveform, we generated a distance dependent attenuated waveform on neighboring channels. Then, the waveform from its reference channel (where spike showed it largest amplitude) of a spike to recording channel *ch* is attenuated by a Gaussian function:

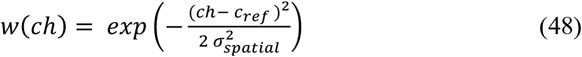

where *c*_ref_ is the reference channel of the spike event and *σ*_spatial_ controls the spatial spread.

For the nonlinear waveform modeling, and incorporating voltage-dependent kinetics and curved spike manifolds, we used each of the canonical waveform above each canonical waveform and transformed it as:

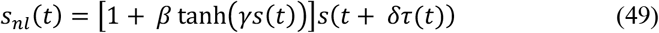

Where 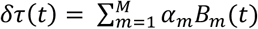, with cubic B-spline basis *B*_*m*_, and coefficient *α*_*m*_*~* 𝒩(0, *σ*_*τ*_),*β* ~ *U (β*_*min*_, *β*_*max*_), and *γ* ~ *U (γ*_*min*_, *γ*_*max*_), which ranges by ≲ 30% curvature and ~ 20–40 μs timing jitter. In the nonlinear modeling, we ranged the number of units showing non-linear waveform variabilities from 10, 25, and 50% of all units (**Figure S5**).

### Modeling spatial drifts

To model electrode drift, the reference channel *c*_ref_ varies over time in one of the following two ways; smooth (sinusoidal) drift:

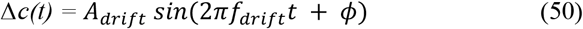

and step drift:

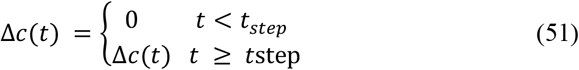

with *c*_ref_ (*t*) = *c* + Δ*c*(*t*). The *f*_drift_ indicates the frequency of the drift, and *ϕ* is the delay phase which is consistent for all units showing drifts. We also considered non-linear stochastic drift for additional simulations shown in **Figure S5** as:

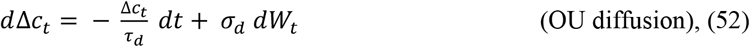

where *τ*_*d*_ is a decay time constant controlling how quickly deviations relax back toward zero, *σ*_*d*_ is the Diffusion coefficient indicating the size of the small, continuous fluctuations, *W*_*t*_ models a small Wiener process. The final drift model is implemented through adding the Ornstein– Uhlenbeck diffusion process as a nonlinear stochastic component to the continuous sinusoidal drift as:

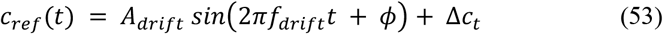

### Multichannel signal

After generating both spike trains and background LFP activity for each channel, we generated the multichannel signal containing both. The time-domain voltage on channel *ch* at time *t* is the sum of LFP noise and spike contributions and is then calculated as:

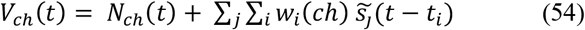

where the sum runs over all spike events of all unit *s*_j_, occurring at times *t*^*i* 1,36^.

### Benchmarking metrics

To evaluate spike-sorting performance, we matched the detected clusters to ground-truth classes and computed standard metrics. For *K* detected cluster 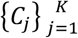 and *M* ground-truth classes 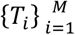, we assigned each cluster to the class by minimizing the Manhattan distance:

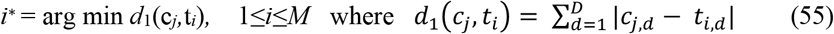

For each true class *T*_*i*_, we computed precision (P_i_), recall (R_i_), F1-score (F_1,i_), and accuracy (A_i_):

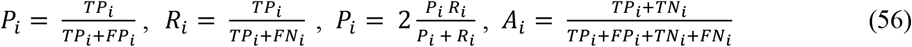

where *TP*_*i*_, *FP*_*i*_, *TN*_*i*_, *FN*_*i*_ denote true positives, false positives, true negatives, and false negatives for class i, respectively. Under each simulation condition, these metrics are calculated for all classes and compared between KIASORT and Kilosort4 using the Wilcoxon signed-rank test.

## Acknowledgments

We thank Kari Hoffman, Adam Neumann, and David Brust for their help and feedback with the research. This work was supported by a C.V. Starr Fellowship from Princeton University (KBB), the National Institute of Mental Health (R01MH123687, TW). The funders had no role in study design, analysis, and the decision to publish, or the preparation of this manuscript.

## Author Contributions

K.B.B developed and designed the methodology, modeling, software, analysis. K.B.B wrote the original draft. S.K. and T.W. provided supervision. All authors contributed writing and editing the paper.

## Data and code accessibility

The whole pipeline and code for KIASORT is available from https://kiasort.com.

**Figure S1.**
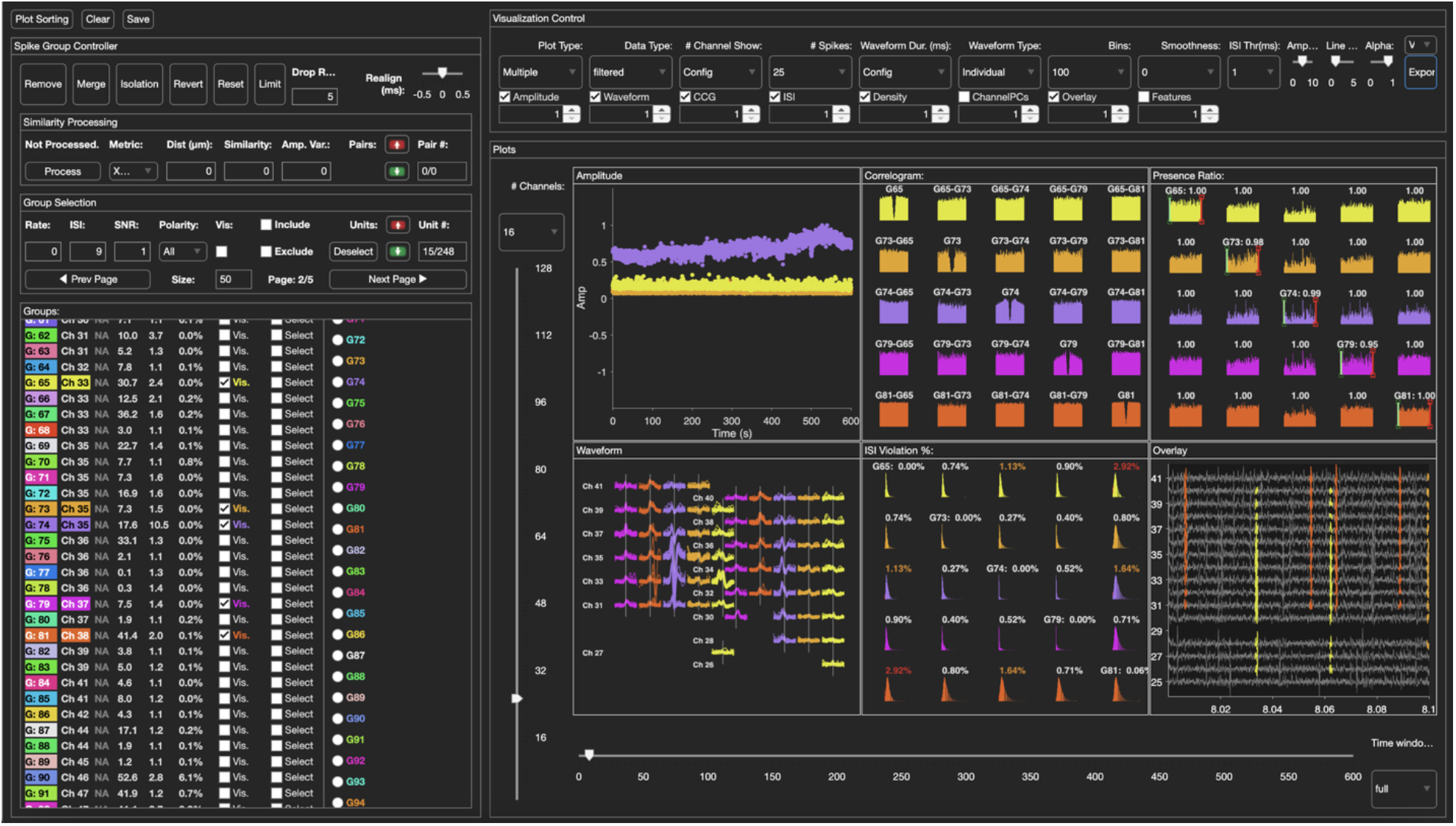
KIASORT Graphical User Interface (GUI). KIASORT includes a GUI that provides a unified platform for data inspection, sorting, and post-hoc curation. A screenshot of the Curation tab which allows users to visualize sorted units; remove or merge units; change isolation types; perform pairwise waveform-similarity processing; and inspect clusters via cross-correlograms, ISI distributions, presence ratios, waveforms, and low-dimensional feature embeddings on each channel.

**Figure S2.**
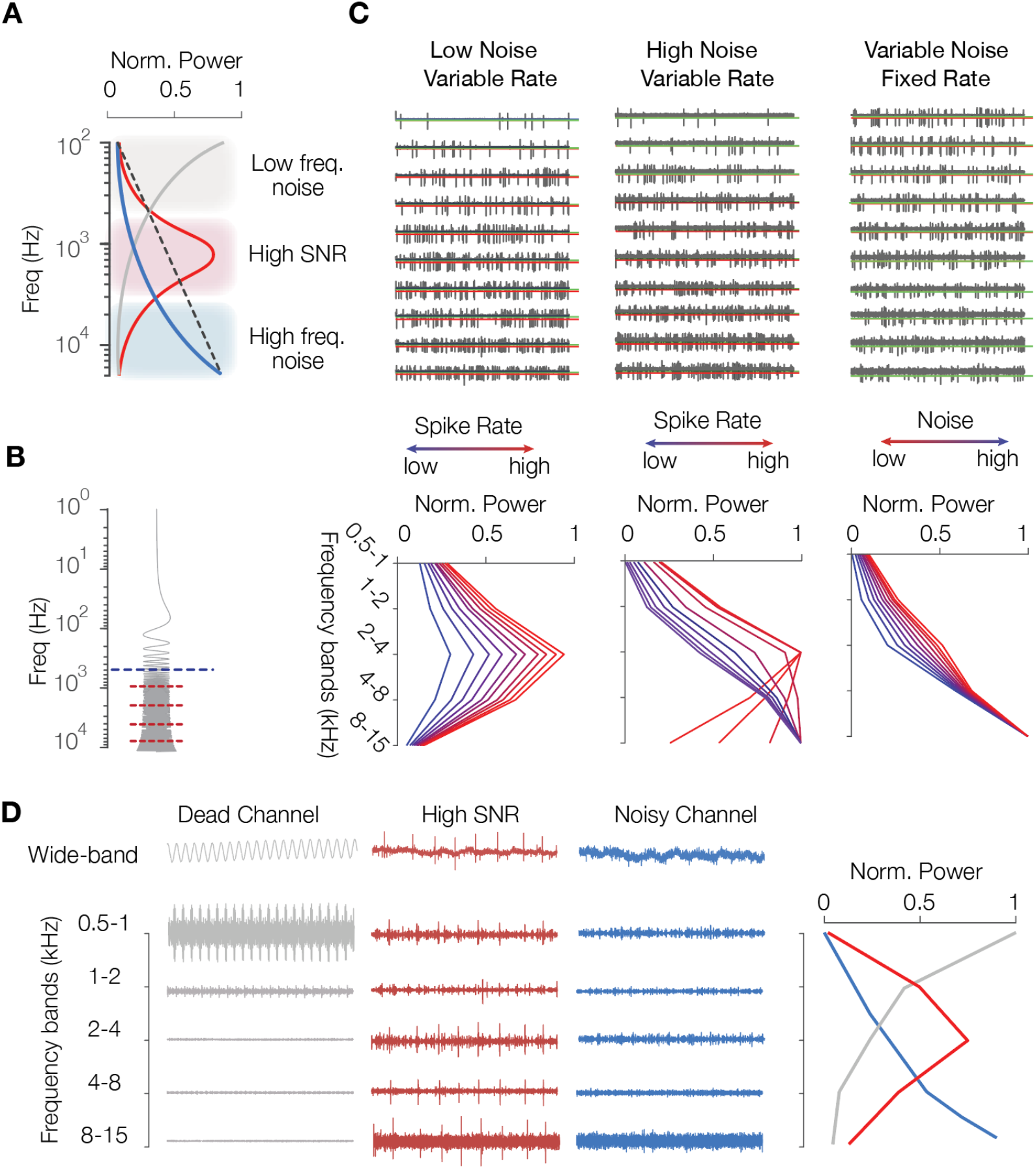
Channel quality check by multiband decomposition. **A**. For an electrode in brain tissue, the dominant signal activity is in the middle frequency band (red), reflecting spiking and other neuronal activity. For a silent or dead channel, the signal is dominated by low-frequency noise (gray), and for a noisy or impaired channel, by high-frequency noise (blue). **B**. To assess different contributing factors to this distribution while remaining computationally efficient, each channel’s signal was band-pass filtered into non-overlapping frequency bands. **C**. Simulated signals generated by varying noise levels and firing rates show that as neural activity increases and noise decreases, the power distribution across frequency bands tends toward a Gaussian shape. **D**. Performance of the algorithm on different channel types from real primate recordings.

**Figure S3.**
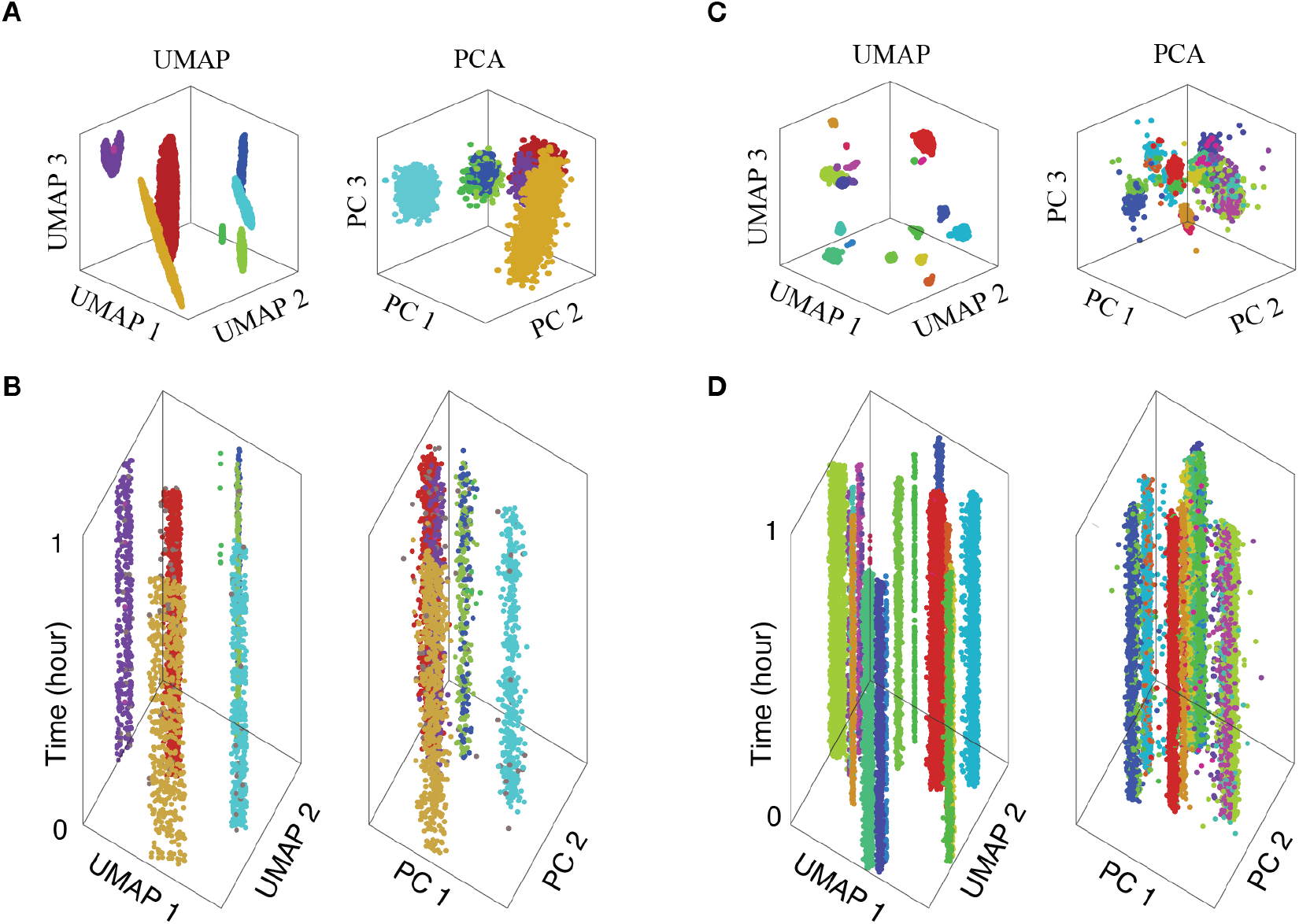
UMAP and PCA feature distributions of clusters. **A, B**. UMAP and PCA embeddings of clusters identified by iterative DBSCAN on a primate laminar probe recording channel: (**A**) 3D embedding, (**B**) 2D embedding overtime colored by cluster id. **C, D**. Same as **A** and **B**, but for a channel from the simulated dataset.

**Figure S4.**
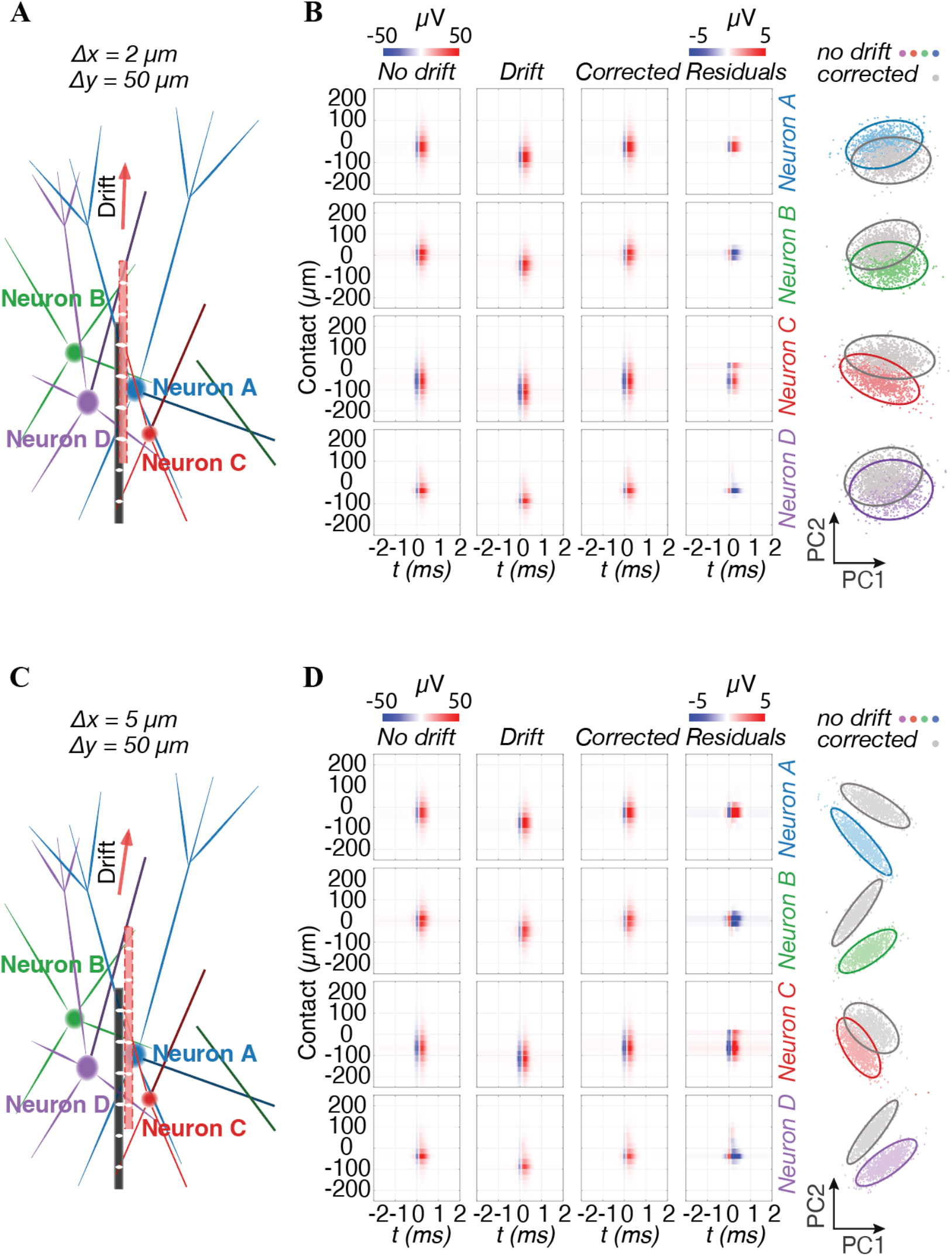
Biophysical model of drift-induced waveform distortions. **A**,**B**: same as in **Figure 4A,B** for Δx = 2 µm. **C**,**D**: same as in **Figure 4A,B** for Δx = 5 µm.

**Figure S5.**
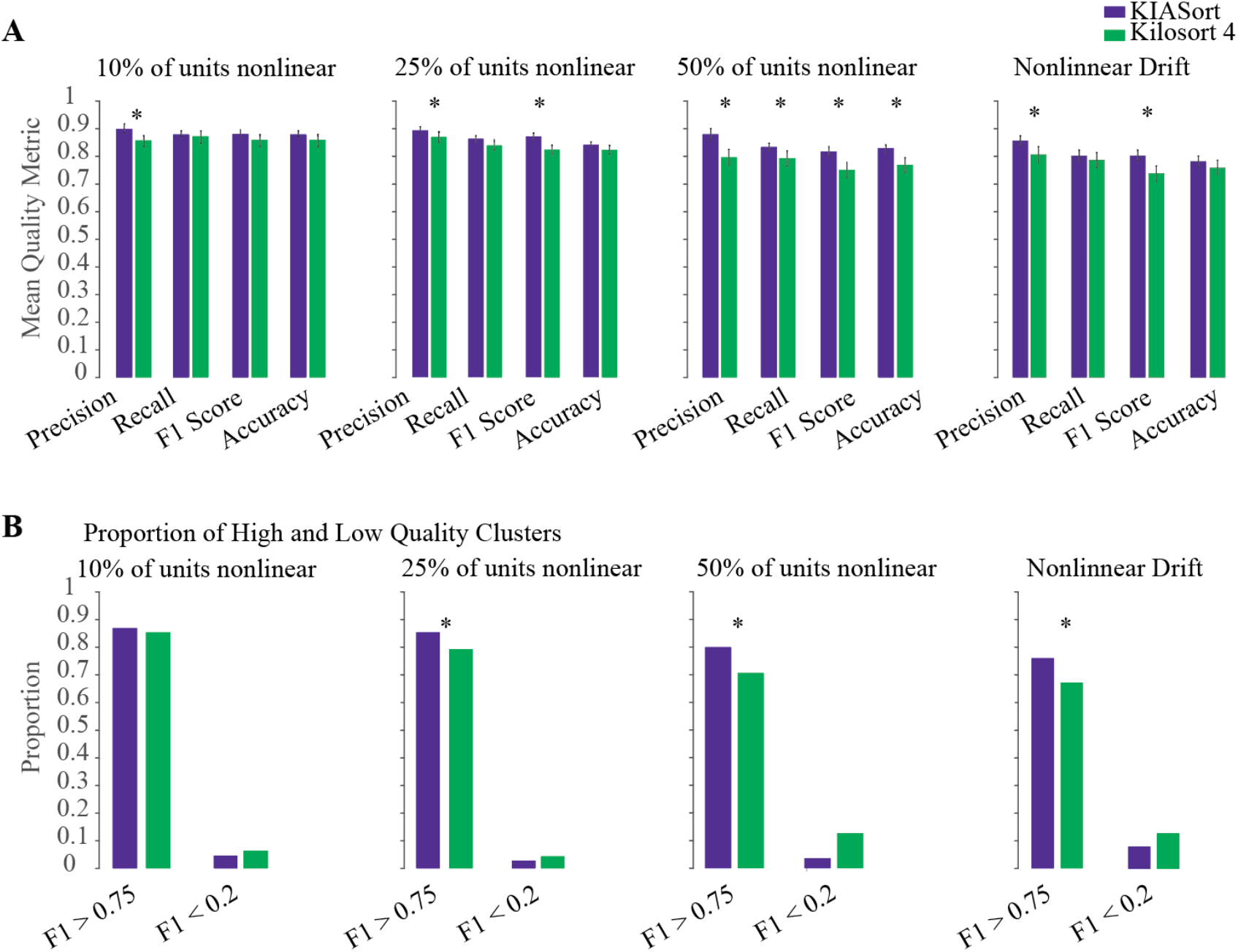
Benchmarking with nonlinear waveform changes and drift. **A, B**. Same as in **Figure 4D,E**, for simulated datasets containing 10%, 25%, and 50% of units showing nonlinear waveform changes, and for simulations implementing nonlinear drift via stochastic per-neuron perturbations combined with sinusoidal probe-based drift.

**Figure S6.**
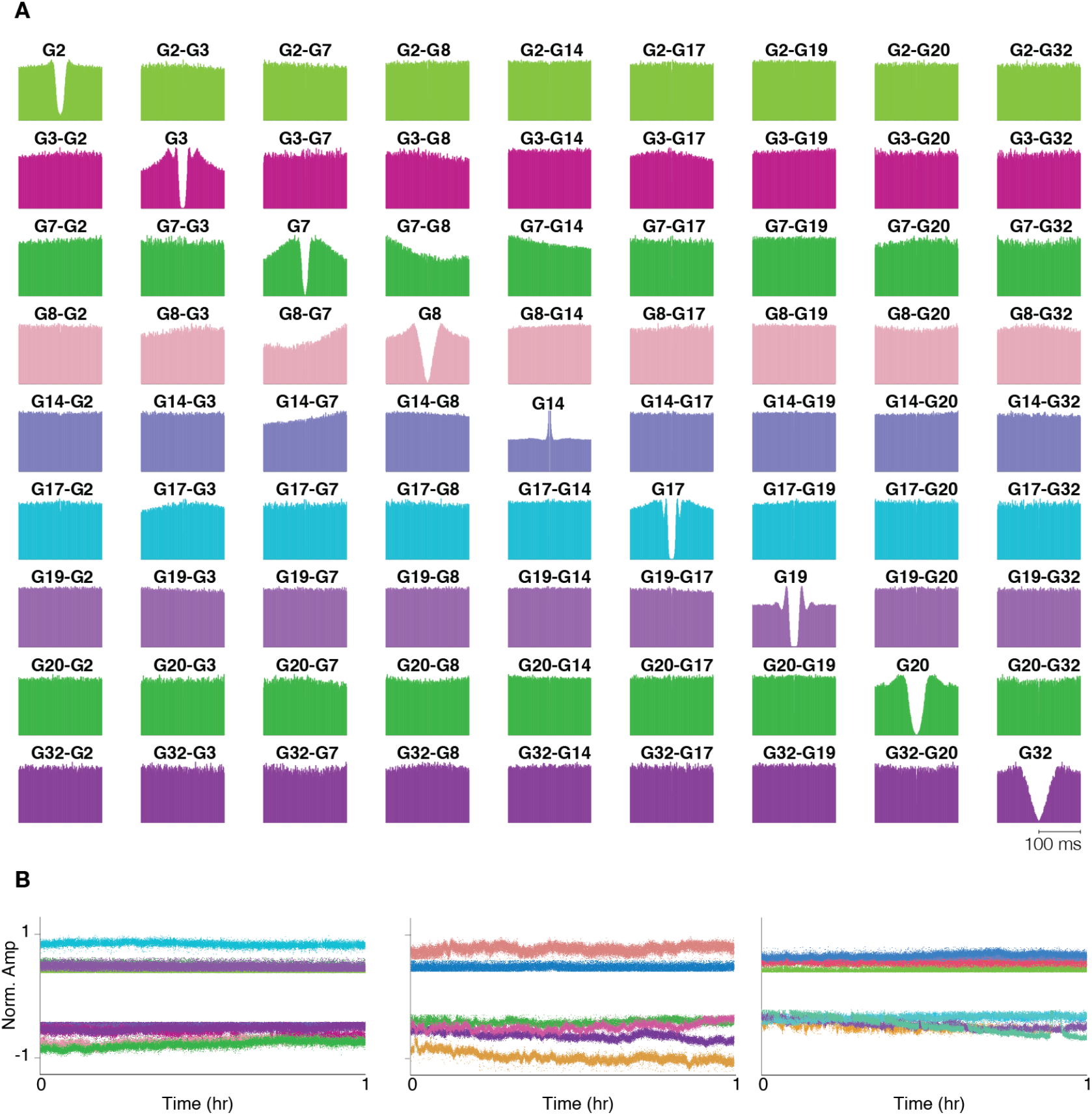
Cross-correlograms and per-neuron drift examples. **A**. Auto- and cross-correlograms (ACG and CCG) between identified neuron groups within a 100 µm radius for different types of units, including fast spiking units. **B**. Three examples of per-neuron drift patterns across three different recording areas in NHP brain: striatum, lateral prefrontal cortex, and anterior cingulate cortex. The left panel corresponds to groups in panel (A). In all three recordings units are selected from a 100 µm radius and show different drift pattern.

